# Altered T cell reactivity to β-amyloid-related antigens in early Alzheimer’s disease

**DOI:** 10.1101/2024.06.04.597424

**Authors:** Chiara Rickenbach, Anna Mallone, Lars Häusle, Larissa Frei, Sarina Seiter, Colin Sparano, Tunahan Kirabali, Kaj Blennow, Henrik Zetterberg, Maria Teresa Ferretti, Luka Kulic, Christoph Hock, Roger M. Nitsch, Valerie Treyer, Anton Gietl, Christoph Gericke

## Abstract

There is growing evidence that the adaptive immune system and neurodegenerative Alzheimer’s disease (AD) are intertwined in multiple ways. Recent studies have reported alterations of the adaptive immune system in early AD stages, such as preclinical AD and mild cognitive impairment (MCI) due to AD. However, the identity of specific antigenic targets and whether the respective response is beneficial or detrimental during disease progression are still open questions. Herein, we describe cross-sectional analyses of blood and cerebrospinal fluid from three different study populations covering early AD stages. We employed high-dimensional mass cytometry, single-cell RNA-sequencing, *in vitro* T cell secretome analysis, and antigen presentation assays to achieve a comprehensive characterization of adaptive immune cell populations. Our results show that subjects at the stage of asymptomatic, preclinical AD can mount a CD4^+^ T helper cell response towards β- amyloid peptide and display an early enrichment of cytotoxic CD8^+^ effector/TEMRA cells in CSF, combined with a less immunosuppressive gene signature of peripheral regulatory T cells. Conversely, in MCI due to AD, we observed increased frequencies of CD8^+^ effector/TEMRA cells in the periphery, characterized by a pro-inflammatory gene expression profile and an overall decrease in antigen responsiveness. Our results demonstrate the complexity of adaptive immune changes in early AD and suggest that it may be beneficial in the preclinical stage to promote specific CD4^+^ T cell responses, while in MCI it may be important to therapeutically target CD8^+^ T cell responses if these prove to be harmful.

## Introduction

Alzheimer’s disease (AD) is a neurodegenerative disease affecting over 50 million people worldwide ^1^. Hallmarks of AD are aggregation and deposition of misfolded β-amyloid peptide (Aβ) into Aβ plaques, neurofibrillary tangles of hyperphosphorylated tau, and neurodegeneration ^2^. After a long asymptomatic phase with Aβ accumulation followed by tau pathology, AD eventually progresses to mild cognitive impairment (MCI) and dementia ^3-7^. Nowadays, biomarkers for AD, such as cerebral Aβ and tau aggregates measured by positron emission tomography (PET) scans and levels of phosphorylated tau species (p-tau) in cerebrospinal fluid (CSF) or plasma, are increased in AD and can predict the probability of a clinical transition from cognitively healthy individuals to MCI and dementia ^8-10^. As particularly the preclinical phase of AD is not well understood, its in-depth characterization would allow improving biomarkers and potentially preventive therapeutic strategies before the onset of symptoms.

In recent years, the role of adaptive immunity in AD has been increasingly recognized. Both clinical and genetic data suggest an involvement of inflammation-related pathways in AD pathogenesis and progression ^11,12^. Multiple studies have identified increased T cell infiltration in AD brains ^13-15^, sparking the question of whether the underlying responses might be beneficial or detrimental during disease progression. For the cytotoxic CD8^+^ subtype of T cells, CD45RA-reactivated T effector memory (TEMRA) cells were shown to be increased in peripheral blood and CSF of AD patients ^13^. Interestingly, immunophenotyping studies focusing on earlier stages of AD revealed that CD8^+^ TEMRA cells are altered even before the conversion to AD dementia as early as in preclinical AD ^16^. While lymphocyte infiltration in AD brains appears to be a targeted process, the antigenic target remains unknown ^17^. Some studies investigated the T cell antigen specificity in AD and reported reactivity of CD4^+^ T cells towards Aβ-derived epitopes in a human leukocyte antigen (HLA) class II-dependent manner ^18-20^. These results were no surprise since earlier studies observed that immunization with Aβ peptide resulted in less AD pathology in mouse models of β-amyloidosis ^21^. The subsequent first active immunization strategy for AD was AN-1792, mainly consisting of synthetic full-length Aβ1-42 peptide. Clinical trials were suspended due to cases of meningoencephalitis ^22^. Nevertheless, T cell recruitment to the CNS occurred, and some responders mounted antibody responses against the amino-terminus (N-terminus) of Aβ ^23^. Further discoveries of human autoantibodies directed against aggregated Aβ support the concept of successful priming of the adaptive immune system in AD ^24-26^.

To investigate the onset, targets, and nature of AD-associated T cell responses, we performed an in-depth cross-sectional immunophenotyping of three study populations covering the early stages of AD. We included cognitively healthy subjects with early evidence of cerebral β-amyloidosis (referred to as preclinical AD) and subjects with mild cognitive impairment (MCI) due to AD (also referred to as prodromal AD). We provide a detailed analysis of immune cell populations in peripheral blood and CSF by combining mass cytometry by time-of-flight (CyTOF) and single-cell RNA sequencing (scRNA-seq). Additionally, we performed a functional analysis of the T cell secretome and antigen presentation assays. We report that CD4^+^ T cells in preclinical AD display a type 2 T helper profile and reactivity against carboxyl-terminal (C-terminal) and mid-sequence peptides derived from linear Aβ1-42. In contrast, we observe an increased frequency and a more pro-inflammatory gene expression of CD8^+^ TEMRA/effector cells in MCI subjects. Our analysis suggests a potentially beneficial role of activated CD4^+^ T helper cells in preclinical AD and a potentially harmful involvement of cytotoxic CD8^+^ T cells in later stages of AD.

## Methods

### Study Participants

We used peripheral blood mononuclear cells (PBMCs), blood plasma, and serum from study participants divided into healthy control subjects (HCS) and participants diagnosed with mild cognitive impairment (MCI, according to consensus criteria ^27^). All participants are part of an ongoing prospective longitudinal cohort study with extensive multimodal imaging, clinical and cognitive assessments (see ^16,28^ for detailed characteristics on ‘ID-Cog’ study, as well as **Table S1-3**). For the analyses, three subpopulations of study participants were selected from the overall study population. Participants from study population 1 represent a subgroup of participants who consented to lumbar puncture for CSF collection and were grouped based on the ratio of AD biomarkers Aβ42 and Aβ40 in the CSF. Participants from study populations 2 and 3 were assessed for cerebral Aβ load measured via positron emission tomography (PET) imaging using [18F]-Flutemetamol (FMM) tracer. The median time difference between PBMC sampling and PET imaging was 19 days. To standardize quantitative Aβ imaging measurements, the ‘Centiloid’ approach was used ^29^. The Centiloid cut-off for Aβ positivity was set at FMM Centiloid > 12, which has been shown to mark the transition from absence of pathology to subtle pathology ^30^. Study participants were grouped based on Aβ PET into Aβ-negative and Aβ-positive HCS (HCS- and HCS+), and Aβ-positive MCI (MCI+ = MCI due to AD). Study populations 2 and 3 additionally underwent AD biomarker analysis for Aβ42 and Aβ40, as well as phospho-tau 217 (p-tau217), neurofilament light (NfL) and glial fibrillary acidic protein (GFAP) in blood plasma. Subjects with current/recent tumors, Hashimoto’s disease or increased serum levels of C-reactive protein (CRP) were excluded.

### APOE Genotyping

*APOE* genotyping was performed using a restriction isotyping protocol ^31^ or Sanger Sequencing (Microsynth AG).

### PET Imaging and Image Analysis

Standard reconstructed PET/MR [18F]-FMM-PET images were averaged over the time window between 85-105 minutes post injection. Images were acquired on a Signa PET/MR (GE Healthcare). Participants received a maximum of 140 MBq of tracer. Images were processed with PMOD 4.0 NeuroTool (PMOD Technologies, Zurich, Switzerland) according to standard Centiloid methods ^29^.

### Neuropsychological Testing

Details of the neuropsychological testing within the cohort study were published previously ^16,28,32^. For this analysis, we used Mini-Mental State Examination (MMSE) assessment for global estimate of cognitive functions.

### Serology for Virus Antibody Titers

Viral antibody titers were analysed from serum samples via ELISA or indirect immunofluorescence at the Institute of Medical Virology, University of Zurich. Antibody titers against Cytomegalovirus (CMV) (IgG and IgM), Epstein-Barr virus (EBV) (IgG and IgM against viral capsid antigen (VCA) and IgG against Epstein-Barr nuclear antigen (EBNA)), Herpes simplex virus (HSV) 1 and 2 (IgG and IgM), and Varicella-Zoster-Virus (VZV) (IgG and IgM) were measured. Quantitative results were obtained for the following antibody titers: VZV IgG (in mIU/ml, threshold for positive test >100), CMV IgG (in AE/ml, threshold ≥6), HSV-1 IgG (in ‘Signal divided by cut-off’: S/CO, threshold >1) and HSV-2 IgG (in S/CO, threshold >1). All other titers were analyzed qualitatively (positive/negative).

### Fluid AD Biomarkers

Blood of study participants was collected in ethylenediaminetetraacetic acid (EDTA) tubes (Vacutainer EDTA Tubes, BD), inverted 10 times, and centrifuged at 1’620 x g for 12 min at 6°C. Plasma supernatant was aliquoted and stored at -80°C. CSF samples were centrifuged max. 30min after lumbar puncture at 350 x g for 10min to separate CSF cells from supernatant. CSF supernatant was aliquoted and frozen at -80°C. For biomarker analysis of study population 1, quantifications were performed by Quanterix corporation on a Simoa HD-X instrument using Simoa Human Neurology 3-plex A (N3PA) immunoassays for measuring CSF Aβ42, Aβ40 and total tau. Biomarker analysis for study populations 2 and 3 was performed in the laboratories of Prof. H. Zetterberg and Prof. K. Blennow (Sahlgrenska University Hospital, Mölndal, Sweden). In brief, quantifications of plasma markers Aβ42, Aβ40, NfL, and GFAP were performed with a customized Neurology 4-plex A platform (Quanterix), analysis of plasma p-tau217 was measured with an in-house assay of University of Gothenburg. Fluid biomarker levels below the limit of quantitation (LOQ) were set at LOQ/2 according to standard approaches for left-censored LOQ data ^33^.

### C-reactive Protein (CRP) Assessment

Serum levels of CRP were analysed as part of routine diagnostics in externally certified laboratories to exclude acute infections and inflammation.

### Staining for Mass Cytometry

Cryopreserved PBMCs were thawed at 37°C and washed with cell culture medium (RPMI 1640 (Sigma) with 10% v/v heat-inactivated FBS (Gibco) and 1% v/v Glutamax (Gibco)). 1 million PBMCs from each subject were transferred to a 96-well plate, washed in PBS and incubated with Cell-ID ^198^Pt-Cisplatin (Fluidigm) 1:10’000 in PBS for 10min at room temperature (RT) for dead-cell exclusion staining. Cells were then incubated with Fc receptor blocking antibodies (Human TruStain FcX, Biolegend) 1:20 in Cell Staining Buffer (CSB, Fluidigm) for 5 minutes at RT, followed by barcording for 20min at RT. Customized live-cell barcoding was used, based on combinations of anti-human CD45 antibodies labelled with different metal tags (10-choose-4 approach for a total of 210 possible combinations).

Cells were washed in CSB, pooled and incubated with a CyTOF antibody panel (for 20min at RT) including classical lineage markers for PBMCs and specific T cell activation markers (**Table S6**). After washing in CSB, cells were fixed with 1.6% v/v formaldehyde in CSB and stored overnight at 4°C. Stained samples were aliquoted in 20 million cell batches and frozen in 10% v/v DMSO in FBS for long-term storage at -80°C. Frozen samples were thawed in pre-warmed PBS and incubated in Cell-ID Intercalator-Ir (Fluidigm) 1:5000 in Maxpar Fix and Perm Buffer (Fluidigm) for 1 hour at RT. After washing in deionized metal-free water and in Cell Acquisition Solution (CAS, Fluidigm), samples were diluted to 1.2 million cells/ml in CAS containing 1:10 EQ Four Element Calibration Beads (Fluidigm). Samples were analyzed with a Helios CyTOF2 mass cytometer (DVS Sciences, Fluidigm).

### Isolation of CSF Cells and Staining for Mass Cytometry

Lumbar puncture with an atraumatic needle was used to collect 6ml of CSF from each study participant. Within 30min after lumbar puncture, CSF samples were centrifuged at 350 x g for 10min to separate CSF cells from supernatant. CSF supernatant was aliquoted and frozen for further biomarker analysis. CSF cells were incubated with Cell-ID ^198^Pt-Cisplatin (Fluidigm) 1:10’000 in PBS for 10min at RT for dead-cell exclusion staining. ^198^Pt-Cisplatin staining was quenched with cell culture medium. CSF cells were fixed in 1.6% v/v formaldehyde for 1 hour prior to freezing in 10% v/v DMSO in FBS to increase recovery yield after thawing.

Cryopreserved CSF cells were thawed at 37°C and washed with cell culture medium. Due to low CSF cell numbers (<5’000 cells/ml) and to reduce CSF cell loss during subsequent staining steps, CSF cells were mixed with rat PBMC dump cells (1 Mio. cells per CSF cell sample, Creative Biolabs) pretreated with Rat Fc Block™ Reagent (BD Biosciences). Further Fc receptor blocking, barcoding and staining with a CyTOF antibody panel (**Table S6**) were performed as described above for PBMCs.

### Mass Cytometry Data Pre-processing

The analysis was carried out using an R environment implemented in RStudio. The packages *Premessa* and *CATALYST* were used for batch normalization, file concatenation, debarcoding and compensation to correct for overspilling high-intensity barcoding channels^34-36^.

### Mass Cytometry Data Analysis

Clustering, UMAP and heatmap visualizations were performed with the *CATALYST* package^35^. Clustering was performed with FlowSOM algorithm followed by ConsensusClusterPlus metaclustering ^35^. Landmark clusters were obtained after manual merging based on estimations of expectable subclusters ^16^. To obtain high-resolution unsupervised clustering nodes we used the diffcyt computational framework from the *diffcyt* package ^37^. Cell-type (lineage) markers were used for high-resolution FlowSOM clustering to obtain ‘cluster nodes’ that were projected onto a scaffold map using landmark nodes as reference. Cell-state markers were used for differential state (DS) testing using the ‘diffcyt-DS-limma’ method. A DS node was defined as unsupervised clustering node with a significant cell state marker alteration.

Scaffold maps were constructed to generate a force-directed graph using the *vite* package ^38^. The graph was visualized using Fruchterman-Reingold layout with *igraph* and *ggraph* packages ^39^. Cohen’s d effect size was calculated using *rstatix* package ^40^. Other plots were generated using ggplot2.

### Cell Processing for Single-cell RNA-sequencing

Cryopreserved PBMCs were thawed at 37°C and washed with cell culture medium (RPMI 1640 (Sigma) with 10% v/v heat-inactivated FBS (Gibco) and 1% v/v Glutamax (Gibco)). Single-cell suspensions containing 10’000 cells/subject were analysed. Transcriptomes and TCRs were processed using the Chromium Next GEM Single Cell V(D)J Reagent Kits v1.1 (10X Genomics) according to manufacturer’s instructions. Pre-processing of sequencing data was performed with the Cell Ranger pipeline (10X Genomics) according to manufacturer’s instructions.

### Single-cell RNA-sequencing Data Analysis

scRNA-seq analysis was performed using packages available on Bioconductor ^41^. Briefly, low quality cells were identified and filtered from each sample using the median absolute deviation (MAD) as threshold. Cells with high percentage of mitochondrial genes, with too low/high library size and with too low/high number of genes were excluded.

Normalized expression values for each cell were obtained by library size normalization and log-transformation. The top 10% most highly variable genes were selected for principal component analysis (PCA), and the Harmony algorithm ^42^ was used to harmonize the data from every subject. The function ‘scDblFinder’ from *scDblFinder* package was used to detect and filter doublets ^43^. The Harmony-corrected dataset was used as input for UMAP visualization. For unsupervised graph-based clustering, we used ‘buildSNNGraph’ followed by ‘cluster_louvain’ from *igraph* package. Clusters were manually annotated based on median gene expression profiles. Gene set enrichment analysis (GSEA) was performed with *escape* package. The Enrichment Score (ES) was calculated using the function ‘enrichIt’ of the ‘UCell’ algorithm ^44^ by comparing the cumulative distribution of a gene set’s expression values in the cell cluster of interest with the cumulative distribution of the same gene set in a reference group (other cells in the data set). The normalized ES was compared across the diagnostic groups with a non-parametric Kruskal-Wallis test. The following gene sets were analysed: HALLMARK_TNFA_SIGNALING_VIA_NFKB (including TNF and TNF-response genes), HALLMARK_IL6_JAK_STAT3_SIGNALING (including IL-6 response genes), HALLMARK_INFLAMMATORY_RESPONSE (including CD69, TNF response genes, IL-4 and IL-6 response genes), HALLMARK_IL2_STAT5_SIGNALING (including IL-2 response genes), HALLMARK_IL6_JAK_STAT3_SIGNALING, an anergy gene set (including NR4A3, EGR1, NR4A2, PFKP, GCH1, see reference ^44^), and a cytotoxicity gene set (including NKG7, KLRD1, KLRB1, KLRC1, GNLY, GZMB, NCAM1, among others; see reference ^45,46^). Genes expressed in less than 5% of the cells were excluded.

### *Ex vivo* Reactivation of PBMCs

Cryopreserved PBMCs were thawed at 37°C and washed with cell culture medium (RPMI 1640, Gibco) with 10% v/v heat-inactivated FBS (Gibco) and 1% v/v Glutamax (Gibco). Cells were re-activated *ex vivo* with Cell Activation Cocktail containing Brefeldin A (Biolegend) for 4h at 37°C.

### Antigen-presentation Assays

Cryopreserved PBMCs were thawed at 37°C and washed with cell culture medium (RPMI 1640 (Sigma) with 5% v/v heat-inactivated human serum (Sigma), 1% v/v Glutamax (Gibco), 1% v/v Sodium Pyruvate (Gibco), and 1% v/v non-essential amino acids (Gibco)). In an optimized setup for CD4^+^ T cells, the cells were incubated with 1ug/ml Ultra-LEAF^TM^ purified anti-CD28 (clone CD28.2, Biolegend) agonistic antibody, as well as peptide pools (**Table S4 and S5**) at a concentration of 4ug/ml for 16h at 37°C, followed by *ex vivo* reactivation as described above.

### Surface and Intracellular Cytokine Staining by Spectral Flow Cytometry

Cells were washed in HBSS and incubated in 1:1000 Fixable LiveDead Blue mix. Cells were then incubated with Fc receptor and Monocyte blocking antibodies 1:20 in FACS buffer (HBSS, 2% v/v heat-inactivated FBS, 5mM EDTA, 0.01% v/v sodium azide), followed by incubation with a fluorochrome-conjugated antibody mix (**Table S7**) in FACS buffer for 45min at 4°C. For intracellular cytokine and cytolytic molecule detection, cells were fixed with Cytofix/Cytoperm (BD Biosciences) for 30min at 4°C. Cells were incubated with intracellular antibody mix in permeabilization buffer overnight at 4°C. Samples were acquired with a 5L Cytek Aurora Spectral Analyzer (Cytek Bioscience). The compensation matrix was corrected in FlowJo (TreeStar).

### Flow Cytometry Data Analysis

For secretome analysis, samples were gated for live CD3^+^ singlets in FlowJo (TreeStar) and exported to RStudio. Clustering, UMAP and heatmap visualizations were performed with the *CATALYST* package ^35^. Clustering was performed with FlowSOM algorithm followed by ConsensusClusterPlus metaclustering. A first round of clustering was performed using lineage (‘cell type’) markers as input. To obtain functional clusters, we used functional (‘cell state’) markers. For antigen presentation analysis, cell populations were obtained via manual gating in FlowJo (TreeStar).

### Preparation of Aβ1-42 Oligomers and Fibrils

Aβ1-42 peptide (DAEFRHDSGYEVHHQKLVFFAEDVGSNKGAIIGLMVGGVVIA, rPeptide) was reconstituted in hexafluoroisopropanol (HFIP, Sigma) at a concentration of 5mg/ml, aliquoted, re-lyophilized and stored at -80°C. Aβ1-42 aliquots were reconstituted in DMSO (Gibco, Thermo Scientific) at 5mM, sonicated for 10min and diluted to 100μM in PBS (Gibco, Thermo Scientific). Aβ1-42 oligomers were obtained by orbital shaking for 24h at 4°C; Aβ1-42 fibrils were obtained by orbital shaking for 3 to 6 days at 37°C. All protocols were adapted according to established protocols ^47^. Aβ1-42 oligomeric and fibrillary states were confirmed via SDS PAGE and western blotting.

### Western Blotting

Briefly, 5μg of protein were boiled for 2min at 85°C in Novex^TM^ Tricine SDS Sample Buffer 2X (Invitrogen), separated on Novex^TM^ 10%-20% Tricine Protein Gels (Invitrogen) and transferred on a 0.2μm nitrocellulose membrane (Invitrogen). Blots were blocked in TBS containing 0.05% v/v Tween-20 and 5% w/v milk powder, and incubated overnight at 4°C with mouse anti-Aβ1-16 antibody (Biolegend, clone 6E10). Secondary antibody incubation was carried out for 2 hours at RT using goat anti-mouse IgG H+L – HRP (Jackson). Protein bands were detected with ImageQuant LAS 4000 (GE Healthcare Life Sciences) using ECL Prime Western Blotting Detection Reagent (GE Healthcare).

### Limited Proteolysis (LiP)

After aggregation of 100μM Aβ1-42 peptide into fibrils for 4 days at 37°C on orbital shakers, non-aggregated fragments were separated with ultra-centrifugation at 100’000 x g at 4°C for 60min. Aβ1-42 fibrils were spiked with equal mass of BSA (Sigma) and digested with Proteinase K (Sigma) in 20mM HEPES, 150mM KCl, 10mM MgCl_2_ in PBS (Gibco, Thermo Scientific) for 5min at 25°C ^48^. Digestion was stopped via heat-inactivation at 99°C for 5min and addition of 10% v/v sodium deoxycholate (Sigma) at a final concentration of 5%. Low- and high-molecular weight fragments were separated via ultra-centrifugation at 100’000 x g at 4°C for 60min. Samples were stored at -80°C. Protein digestion was confirmed via silver staining. Liquid chromatography with tandem mass spectrometry (LC-MS/MS) was performed in the laboratory of Prof. Paola Picotti (Institute of Molecular Systems Biology, ETH Zurich).

### Silver Staining

Following boiling and separation of 3μg of protein via SDS PAGE as described above, silver staining was performed using Pierce^TM^ Silver Stain Kit (Thermo Scientific) according to manufacturer’s instructions. Briefly, gels were fixed for 30min in 30% v/v ethanol and 10% v/v acetic acid, followed by washing steps in 10% v/v ethanol and ddH_2_O. Gels were incubated with Sensitizer solution for 1min and with Stain Working Solution for 30min. Developer Working Solution was added for 2-3min until staining appears. The developing reaction was stopped with 5% v/v acetic acid.

### Epitope Prediction via NetMHCIIpan

Epitope prediction was performed with NetMHCIIpan – 4.1 database ^49,50^ with default parameters. 15-mers derived from tau variant 0N4R were screened against HLA haplotypes in the database. Peptides belonging to protein regions predicted to have strong binding scores (within 1% rank) were selected and synthesized (Custom Peptide Synthesis Service, Biomatik) for antigen-presentation assays.

### Statistical Analysis

Statistical analysis was performed in RStudio using *rstatix* and *stats* packages ^40^. Differences between two groups were tested with non-parametric, two-tailed, unpaired Wilcoxon Rank Sum test. Differences between multiple groups were tested with non-parametric Kruskal-Wallis test with correction for multiple comparisons using the Benjamini-Hochberg false discovery rate (FDR) approach. Correlations were calculated with Pearson’s correlation (alpha = 0.05, confidence interval 95%, two-tailed) and non-parametric Spearman’s correlation (alpha = 0.05, confidence interval 95%, two-tailed). Effects of age, sex, and virus titers were estimated using linear regression models, and analysis was performed on the basis of corresponding residuals. Due to small cohort size, an FDR < 10% has been applied. Where possible, violin plots have been used to display numerical data. Horizontal lines represent the median calculated based on the density estimate.

### Ethics Declaration

The participants included in the analyses participated in in-house cohort studies conducted by the ‘Center for Prevention and Dementia Therapy’ at the Institute for Regenerative Medicine at the University of Zurich, Switzerland. All study participants gave written informed consent. The cohort studies including the here shown assessments were approved by the local ethics committee (Kantonale Ethikkommission, Zurich, Switzerland) and conducted in accordance with their guidelines and the Declaration of Helsinki ^51^.

## Results

### Mass Cytometry-based Immunophenotyping Reveals Increased Abundance of Peripheral CD8^+^ TEMRA/Effector Cells in Early AD

To determine the immunophenotypes of individuals in the early AD spectrum, we characterized adaptive immune cells in three study populations, all derived as a subset from a prospective cohort study (**Fig. 1A**). First, we analyzed CSF immune cells from individuals in study population 1, which included cognitively healthy control subjects (HCS) grouped according to CSF Aβ42/40 ratio into brain Aβ-negative (HCS-= high CSF Aβ42/40 ratio = elderly and unaffected) and Aβ-positive (HCS+ = low CSF Aβ42/40 ratio = potentially preclinical AD) (**Fig. 1A-C, Table S1**). Following up on our previous reports ^16^, we investigated whether CD8^+^ TEMRA/effector cells were also increased in the CSF of individuals with early, preclinical AD. After mass cytometry analysis of CSF cells and paired PBMC samples from the same subjects, we applied automated cell clustering using FlowSOM and identified canonical immune cell populations, such as B cells, CD4^+^ and CD8^+^ T cells, myeloid cells (including monocytes and dendritic cells), innate lymphocytes (including NK cells), and unconventional T cells (including double-negative and γδ T cells) (**Fig. S1A-C**). CD4^+^ and CD8^+^ T cell clusters in CSF mainly exhibited an immune profile of memory cells with substantially lower amounts of naïve cells, as expected for CSF-derived cells ^52^ (**Fig. 1D and E, Fig. S1B**). In line with our previous findings, we found that the relative abundance of CD8^+^ TEMRA cells in CSF and in paired blood-derived PBMCs was increased in HCS+ compared to HCS- (**Fig. 1F and G**).

**Fig. 1:**
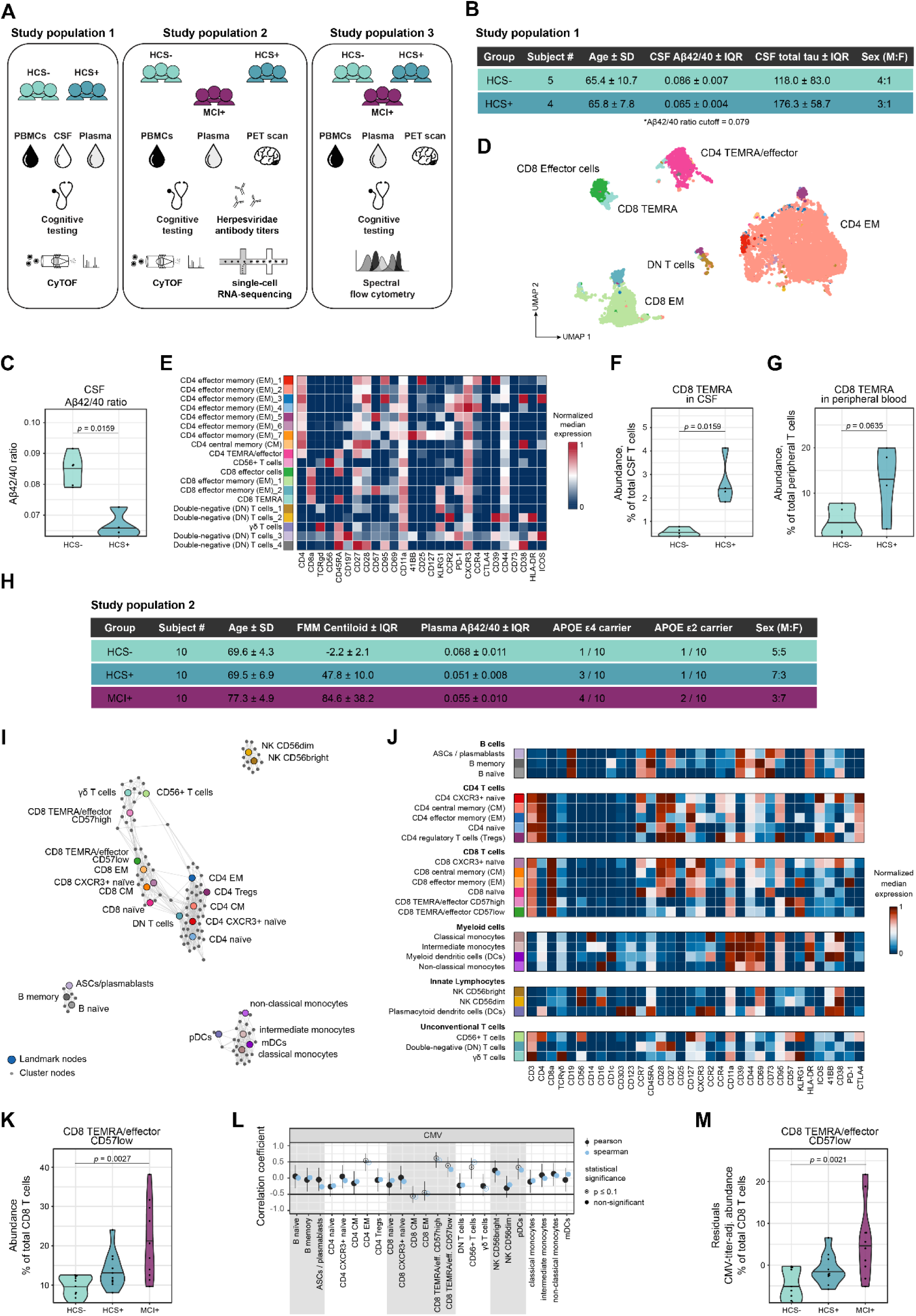
Increased Abundance of CD8^+^ TEMRA/effector Cells in Early AD. (**A**) In total, three study populations were analysed. Study participants were characterized using cognitive testing, Aβ PET imaging, and fluid AD biomarkers. *Study population 1:* Paired PBMCs and CSF immune cells of AD biomarker-negative and -positive cognitively healthy subjects (HCS- and HCS+) were analysed via CyTOF. *Study population 2:* Individuals were divided into cerebral Aβ-negative and -positive cognitively healthy subjects (HCS- and HCS+), and Aβ-positive MCI subjects. PBMCs were analysed via CyTOF and scRNA-seq. Serum was used for quantification of anti-*Herpesviridae* antibody titers. *Study population 3:* Study participants were divided into HCS-, HCS+, and MCI+ groups. Functional analysis of T cell secretome and epitope specificity was performed via spectral flow cytometry. (**B**) Demographic and clinical characteristics of study population 1. (**C**) Violin plot depicting CSF Aβ42/40 ratio in cognitively healthy subjects of study population 1. Subjects were categorized as biomarker-positive if Aβ42/40 ratio in CSF was below threshold value (HCS+, cutoff = 0.079). (**D**) UMAP coloured by CSF T cell subsets obtained with FlowSOM clustering for all non-naïve CD3^+^ cells in the CSF. (**E**) Corresponding heatmap depicting median scaled marker expression within CSF T cell subsets. (**F**) Violin plot showing relative abundance of CD8^+^ TEMRA cells among total CSF T cells. (**G**) Violin plot showing relative abundance of CD8^+^ TEMRA cells among total peripheral T cells. (**H**) Demographic, clinical, and genetic characteristics of study population 2. Subjects were considered cerebral Aβ-positive with FMM Centiloid > 12. (**I**) Scaffold map of total CD45^+^ cells analyzed via CyTOF. Each node represents a cell cluster, including landmark nodes (manually annotated and merged FlowSOM clusters, shown in colours) and cluster nodes (unsupervised FlowSOM clusters, in grey). (**J**) Corresponding heatmap depicting median scaled marker expression within each landmark node. (**K**) Violin plot showing relative landmark node abundance for the CD8^+^ TEMRA/effector CD57low node among total CD8^+^ T cells. (**L**) Correlations between CMV IgG titers and relative landmark node abundance. The estimated Pearson’s r correlation coefficients are shown in black, lines correspond to 95% confidence intervals; the estimated Spearman’s rho correlation coefficients are shown in blue. Empty circles indicate *p* values ≤ 0.1. (**M**) Violin plot showing anti-CMV IgG-adjusted landmark node abundance for the CD8^+^ TEMRA/effector CD57low node among total CD8^+^ T cells. Statistical significance was calculated using non-parametric unpaired Wilcoxon rank-sum test (**C, F, G**) or Kruskal-Wallis test with false-discovery-rate (FDR) method of Benjamini and Hochberg (**K, M**). Selected *p* values ≤ 0.1 (FDR 10%) are displayed.

Next, we analyzed study population 2, which included both cognitively healthy HCS and individuals with MCI. Here, based on Aβ PET imaging data, we established subgroups that were clearly Aβ-negative and strongly Aβ-positive (HCS-, HCS+, MCI+ = MCI due to AD) to study the effect of high Aβ biomarker levels on the immune profile using mass cytometry and scRNA-seq (**Fig. 1A and H, Table S2**). Blood plasma was used for quantification of blood-based AD biomarkers Aβ42, Aβ40 and p-tau217. As per definition, HCS+ and MCI+ had significantly higher cerebral Aβ load compared to HCS-, as shown by increased cerebral Aβ PET levels and simultaneously decreased plasma Aβ42/40 ratio (**Fig. S2A-C**). Plasma-based AD biomarker p-tau217 was only elevated in MCI+ compared to HCS-, correlating with higher cerebral Aβ load (**Fig. S2D and E**). The proportion of individuals with at least one copy of the *APOE* ε4 allele, which is linked to higher AD risk, were higher in Aβ-positive groups compared to HCS- (**Fig. 1H**), with one case of homozygous *APOE* ε4 genotype in HCS+. Across all groups, *APOE* ε4 carriers displayed increased brain Aβ load and decreased plasma Aβ42/40 ratio (**Fig. S2F and G**) but no difference in plasma p-tau217 levels (**Fig. S2H**) compared to non-carriers. *APOE* ε2 carriers were not significantly different in cerebral Aβ load, plasma Aβ42/40 ratio or plasma p-tau217 levels compared to non-carriers (**Fig. S2I-K**).

Using CyTOF immunophenotyping for blood-derived PBMCs, we identified main immune cell populations (**Fig. S1D**) with comparable proportions across diagnostic groups (**Fig. S1E**). We then increased the resolution and identified canonical immune cell populations (here defined as ‘landmark nodes’) including subsets of B cells, CD4^+^ and CD8^+^ T cells, myeloid cells, innate lymphocytes, and unconventional T cells (**Fig. 1I and J**). We compared the proportions of landmark nodes across the three diagnostic groups and observed for CD57low CD8^+^ TEMRA/effector cells an increased relative abundance in MCI+ (*p* = 0.0027) compared to HCS- (**Fig. 1K**).

Generally, proportions of peripheral immune cell subsets could be biased by subclinical levels of viral infections. Moreover, particularly herpesvirus infections have been suggested to be associated with AD pathogenesis ^53^. Therefore, study participants were tested for elevated serum immunoglobulin (IgG and IgM) titers against cytomegalovirus (CMV), Epstein-Barr virus (EBV), Herpes simplex virus (HSV) 1 and 2, and Varicella-Zoster-Virus (VZV). As expected, the majority of individuals were tested positive for IgG antibodies against EBV and VZV, while 40% of study subjects tested positive for anti-CMV IgG (**Fig. S1F**). However, no significant differences in antiviral IgG serum titers between diagnostic groups were observed (**Fig. S1G**). We controlled for an influence of herpesvirus infections on the abundances of canonical immune cell populations by correlating serum *Herpesviridae* IgG titers with landmark node abundances. Most *Herpesviridae* IgG titers had no influence on landmark node abundances (**Fig. S1H**). However, anti-CMV IgG titers appeared to have significant effects on a number of landmark nodes, such as CD4^+^ and CD8^+^ memory T cells as well as on unconventional CD56^+^ T cells (**Fig. 1L**).

Since the proportions of peripheral CD8^+^ TEMRA/effector cells were affected by anti-CMV IgG titers (Spearman correlation: -0.5 < rho < 0.5) (**Fig. 1L**), we tested for CMV IgG-adjusted residuals to exclude that the observed difference was due to CMV reactivation. We confirmed the increase of CD8^+^ TEMRA/effector landmark node abundance in MCI+ (*p* = 0.0021) compared to HCS-, even when corrected for a potential influence of CMV infections on CD8^+^ TEMRA/effector cell abundances (**Fig. 1M**). Frequencies of other T cell subpopulations, including CD4^+^ central memory (CM), CD4^+^ effector memory (EM), regulatory T cells (Tregs), and unconventional T cell subsets, were unchanged across the diagnostic groups after applying the adjustment for CMV IgG titer (**Fig. S1I-M**).

To associate CD8^+^ TEMRA/effector immunophenotype with tau pathology, HCS and MCI subjects were subdivided in an alternative grouping according to the median of p-tau217 plasma levels in study population 2 (= 2.46 pg/ml) instead of using Aβ PET data. For readability, we refer to subjects above the median level as ‘p-tau217+’, emphasizing that this is not a clinical cut-off for tau biomarkers. CD8^+^ TEMRA/effector cell abundance was increased in MCI p-tau217+ compared to HCS p-tau217-, which was also confirmed after adjustment for CMV IgG titers (**Fig. S1N and O**).

Multiple linear regression analysis showed that the effects of age on CD4^+^ and CD8^+^ T cell landmark node abundances were minor, with estimates close to 0 (**Fig. S1P**). In addition, only the proportions of B and NK landmark nodes were affected by sex (**Fig. S1P**).

Collectively, our data suggest an expansion of CD8^+^ TEMRA/effector cells in peripheral blood and CSF of subjects in the early AD stages, independently of latent *Herpesviridae* infections. These immune changes are observable as early as in the preclinical asymptomatic AD stage (HCS+).

### Single-cell Transcriptomics Indicate Less Immunosuppression in Preclinical AD Stage

We performed scRNA-seq for the same set of 30 PBMC samples from study population 2 to delineate the function of T lymphocytes in early AD at the transcriptional level. Overall, the scRNA-seq data analysis revealed comparable PBMC populations as in the previous mass cytometry data, including B cells, CD4^+^ and CD8^+^ T cells, innate lymphocytes, and myeloid cells (**Fig. 2A and B**). We did not find significant differences in relative abundances of main PBMC subsets (**Fig. S3A**). Consistent with CyTOF-based surface characterization (**Fig. 1I and J**), we found CD4^+^ naïve, memory, TEMRA/effector, and Tregs within the CD4^+^ T cell compartment (**Fig. 2C and D**). Interestingly, while the relative abundance of CD4^+^ Tregs was unchanged across diagnostic groups (**Fig. 2E, Fig. S2H**), gene set enrichment analysis (GSEA) on this subset revealed decreased enrichment scores for pro-inflammatory (TNF and IL-6 signalling), immunosuppression-related (TGFβ signalling) and Treg maturation-related (IL-2 and STAT5 signalling) gene modules, and in parallel increased scores for anergy-related gene modules in HCS+ compared to HCS- (**Fig. 2F**). Conversely, MCI+ displayed increased enrichment scores for pro-inflammatory, and Treg maturation-related gene modules compared to HCS- (**Fig. 2F**). The enrichment score for the immunosuppression-related gene module remained reduced compared to HCS- (**Fig. 2F**). In the CD8^+^ T cell compartment, we identified CD8^+^ activated naïve, naïve, memory, and TEMRA/effector cells (**Fig. 2G and H**). Single-cell transcriptomics also confirmed the increase of the CD57low CD8^+^ TEMRA/effector cell cluster in MCI+ compared to HCS- (**Fig. 2I**). For this cluster, GSEA showed decreased scores for pro-inflammatory gene modules (inflammatory response, TNF and IL-6 signalling) in CD57low CD8^+^ TEMRA/effector cells of HCS+ compared to HCS-. In contrast, we observed increased enrichment scores of pro-inflammatory gene modules and simultaneously decreased cytotoxicity-related gene modules in MCI+ compared to HCS- (**Fig. 2J**). As for the mass cytometry data, age and sex differences had only minor influences on landmark cluster abundances, with estimates close to 0 in a multiple linear regression analysis (**Fig. S3B**).

**Fig. 2:**
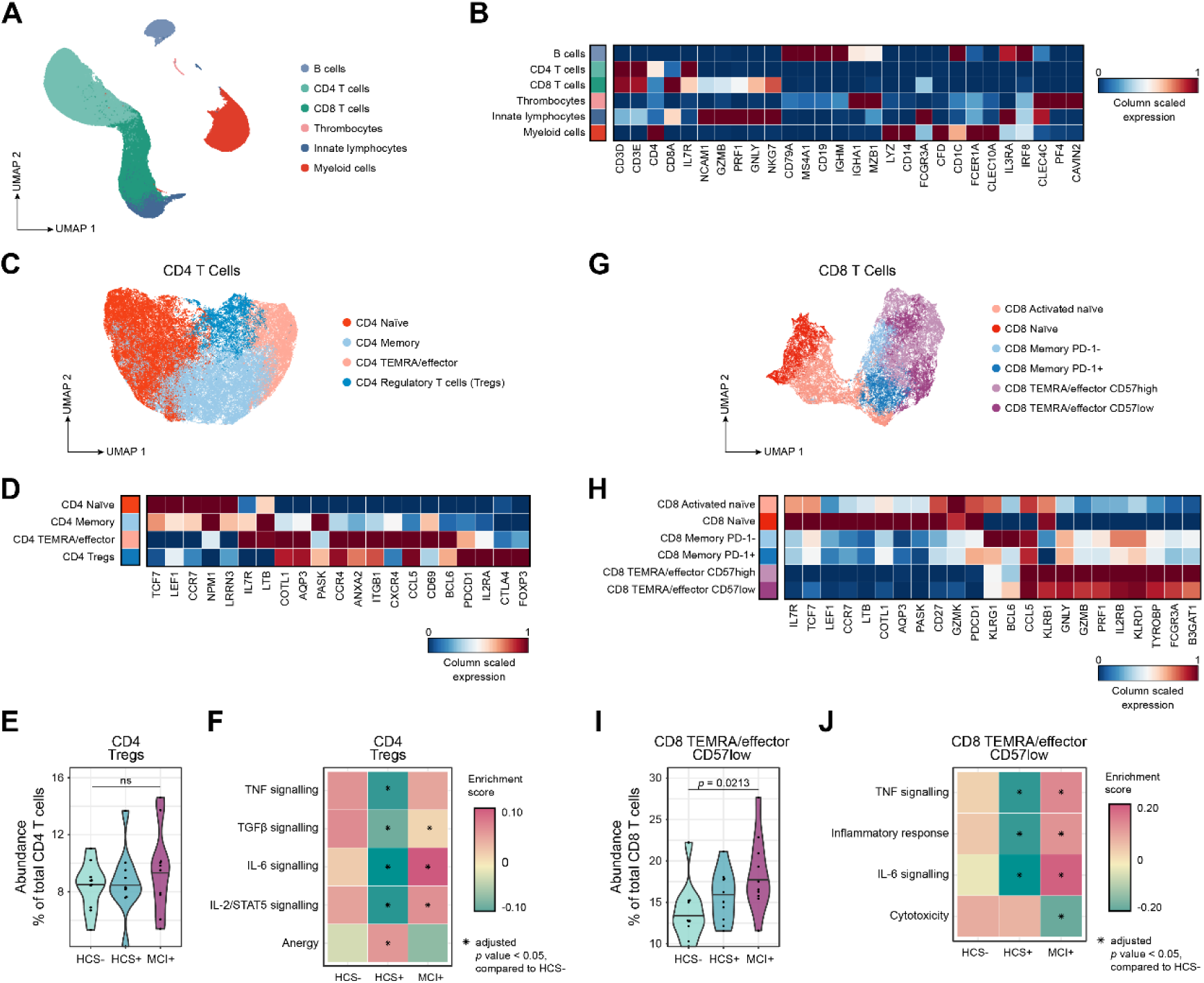
scRNA-seq analysis reveals downregulated CD4^+^ Treg gene signatures in preclinical AD. (**A**) UMAP of total PBMCs in scRNA-seq data, coloured by cluster assignment according to unsupervised graph-based clustering with Louvain algorithm. (**B**) Heatmap depicting scaled gene expression within each PBMC cluster. (**C**) UMAP of CD4^+^ T cell sub-clusters in scRNA-seq data coloured by cluster assignment according to unsupervised graph-based clustering with Louvain algorithm. (**D**) Heatmap depicting scaled gene expression within each CD4^+^ T cell subcluster. (**E**) Violin plot showing relative abundance of CD4^+^ regulatory T cells (Tregs) among total CD4^+^ T cells. (**F**) Heatmap depicting enrichment scores for CD4^+^ Tregs in the 3 diagnostic groups for selected gene sets. (**G**) UMAP of CD8^+^ T cell sub-clusters coloured by cluster assignment according to unsupervised graph-based clustering with Louvain algorithm. (**H**) Heatmap depicting scaled gene expression within each CD8^+^ T cell subcluster. (**I**) Violin plots showing relative abundance of CD8^+^ TEMRA/effector CD57low cluster among total CD8^+^ T cells. (**J**) Heatmap depicting enrichment scores for the CD8^+^ TEMRA/effector CD57low cell cluster in the 3 diagnostic groups for selected gene sets. Statistical significance was calculated using non-parametric Kruskal-Wallis test with false- discovery-rate (FDR) method of Benjamini and Hochberg. Selected *p* values ≤ 0.1 (FDR 10%) are displayed.

Taken together, the single-cell transcriptional profiling of CD4^+^ and CD8^+^ T cell populations in the blood of subjects with early AD indicates a possible impairment of CD4^+^ Treg function and peripheral tolerance in the preclinical AD stage (HCS+). Potentially, this could provide the basis for developing adaptive immune responses against auto-antigens such as Aβ- based aggregates in preclinical AD. On the other hand, in MCI+, a less immunosuppressive Treg and a more pro-inflammatory CD8^+^ TEMRA/effector transcriptomic signature seemingly accompany neurodegeneration and the onset of cognitive decline.

### Altered T cell Reactivity Against Linear Aβ Peptide in Early AD Stages

To investigate the functional component of T cells in the early AD spectrum, we analyzed the T cell secretome and reactivity against AD pathology-related antigen pools. For this, we used PBMCs from study population 3, which includes HCS and MCI subjects grouped based on Aβ PET imaging into cerebral Aβ-negative and Aβ-positive subjects (**Fig. 3A, Table S3**). HCS+ and MCI+ exhibited significantly higher cerebral Aβ load compared to HCS-, as evidenced by increased cerebral Aβ PET levels and concomitantly decreased plasma Aβ42/40 ratio (**Fig. S4A-C**). Plasma-based AD biomarker p-tau217 was elevated in MCI+ compared to HCS, correlating with higher cerebral Aβ load (**Fig. S4D and E**). The proportion of individuals who were hetero- or homozygous for the *APOE* ε4 allele was higher in Aβ- positive groups than in Aβ-negative HCS (**Fig. 3A**), with one case of homozygous *APOE* ε4 genotype in HCS+. *APOE* ε4 carriers had increased brain Aβ load and decreased plasma Aβ42/40 ratio compared to non-carriers (**Fig. S4F and G**). In contrast, plasma p-tau217 levels did not differ between *APOE* ε4 carriers and non-carriers (**Fig. S4H**). *APOE* ε2 carriers were not significantly different in cerebral Aβ load, plasma Aβ42/40 ratio or p-tau217 levels compared to non-carriers (**Fig. S4I-K**).

**Fig. 3:**
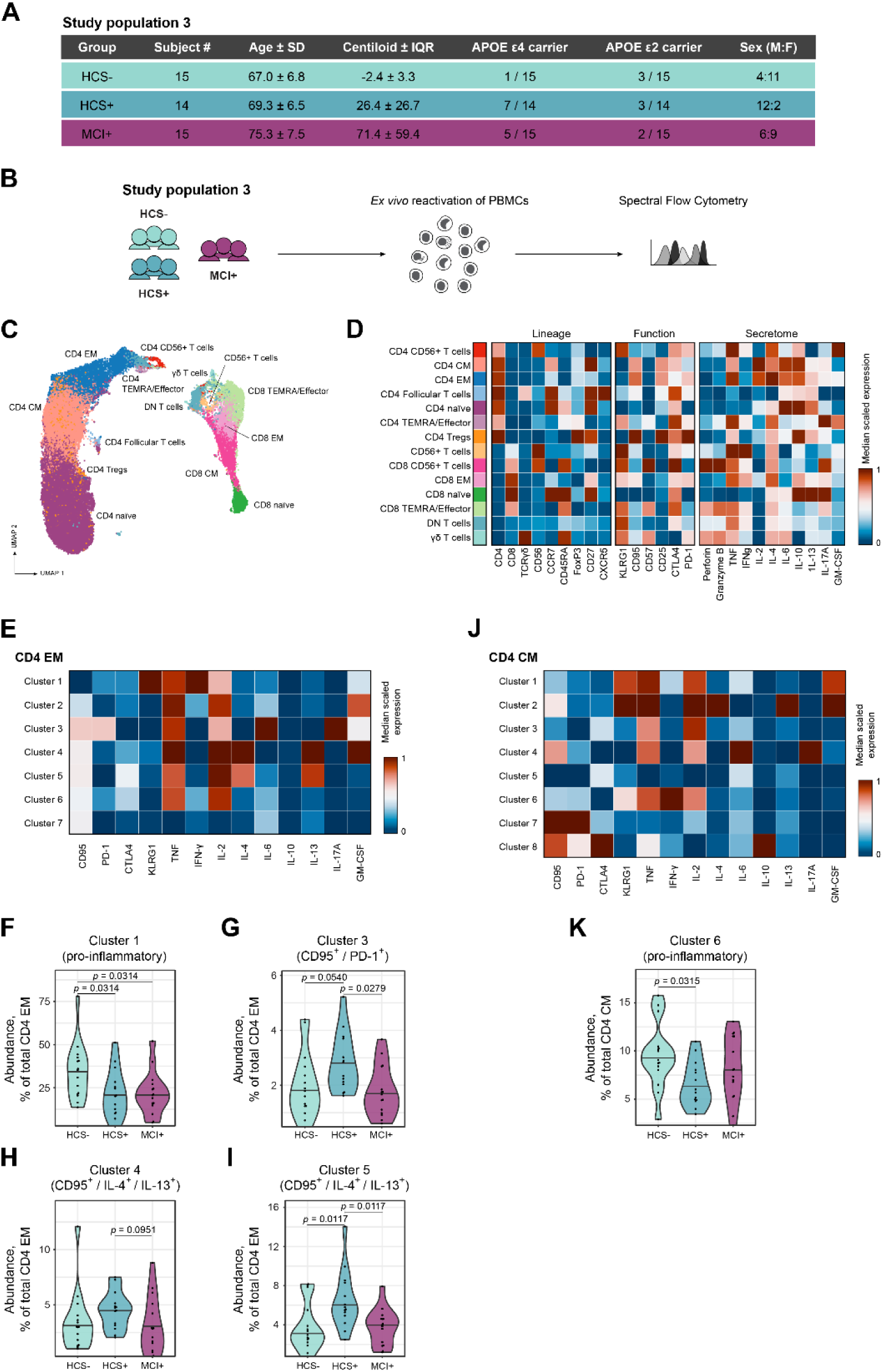
Secretome analysis reveals type 2 T helper-related and anti-inflammatory T cell profiles in preclinical AD. (**A**) Study population 3: Demographic and clinical characteristics. (**B**) PBMCs from study population 3 were reactivated *ex vivo*, T cell secretome was analyzed via spectral flow cytometry. (**C**) UMAP of total T cells coloured by cluster assignment based on unsupervised FlowSOM clustering. (**D**) Corresponding heatmap depicting scaled median expression within each T cell cluster. (**E**) Heatmap depicting scaled median expression of seven functional clusters of CD4^+^ EM obtained via automated FlowSOM clustering using activation markers and cytokines. (**F to I**) Violin plots showing relative abundance of Cluster 1 (**F**), Cluster 3 (**G**), Cluster 4 (**H**), and Cluster 5 (**I**) among total CD4^+^ EM cells. (**J**) Heatmap depicting scaled median expression of eight functional clusters of CD4^+^ CM cells obtained via automated FlowSOM clustering using activation markers and cytokines. (**K**) Violin plot showing relative abundance of Cluster 6 among total CD4^+^ CM. Statistical significance was calculated using non-parametric Kruskal-Wallis test with false-discovery-rate (FDR) method of Benjamini and Hochberg. Selected *p* values ≤ 0.1 (FDR 10%) are displayed.

We first measured the production of cytokines and cytolytic molecules per single cell for *ex vivo* reactivated T cells via spectral flow cytometry (**Fig. 3B**). We identified CD4^+^ and CD8^+^ T cell subsets via automated FlowSOM clustering (**Fig. 3C and D**). Within these subsets, we used activation markers, cytokines and cytolytic molecules to obtain clusters with specific patterns of activation state and secretome. For CD4^+^ EM T cells, we identified seven functional clusters (**Fig. 3E**). A pro-inflammatory CD4^+^ EM T cell cluster containing high levels of Interferon-gamma (IFN-γ) (CD4^+^ EM cluster 1) was decreased in HCS+ and MCI+ groups compared to HCS- (**Fig. 3F**). In contrast, CD4^+^ EM clusters 3, 4, and 5 showed an activation pattern with increased expression of immune homeostasis-related Fas-receptor (CD95) and with higher expression of the immune checkpoint ‘programmed cell death receptor 1’ (PD-1) (CD4^+^ EM cluster 3) or production of type 2 T helper (T_H_2) cell-related cytokines such as IL-4 and IL-13 (CD4^+^ EM clusters 4 and 5). The frequency of these clusters was increased in HCS+ subjects compared to HCS- and MCI+ (**Fig. 3G-I**). For CD4^+^ CM T cells, we identified eight functional clusters, of which CD4^+^ CM cluster 6 displayed a pro-inflammatory immune signature with high levels of IFN-γ and decreased frequency in HCS+ compared to HCS- (**Fig. 3J and K**). For CD8^+^ TEMRA/effector cells, we identified eight functional clusters (**Fig. S5A**). Of these, CD8^+^ TEMRA/effector cluster 8 showed a CD95^+^ activation pattern with high expression of inhibitory immune checkpoint marker CTLA-4, as well as cytokines IL-4, IL-10, and IL-13. This cluster was reduced in MCI+ compared to HCS- and HCS+ (**Fig. S5B**).

For investigating the antigen specificity of T cells, we employed an *in vitro* antigen presentation assay in which we pulsed PBMCs with peptide pools of AD pathology-related putative antigens or positive control antigens derived from influenza, CMV and EBV viruses (**Fig. 4A**). As readout, we measured the surface expression of T cell markers for antigen-specific activation via spectral flow cytometry (**Fig. 4A**). Peptide pools of putative antigen candidates were designed using different approaches including limited proteolysis (LiP) and *in silico* predictions. A previously described source of AD-related peptide antigens is the Aβ1-42 peptide ^19^, the major component of Aβ fibrils and Aβ plaques. To uncover regions of Aβ fibrils that are more prone to structure-dependent digestion and potential presentation to T cells by antigen-presenting cells (APCs), we performed LiP for Aβ fibrils. We produced Aβ fibrils and partially digested them *in vitro* using Proteinase K, a broad-specificity protease that cleaves proteins in a structure-dependent manner ^48^ (**Fig. S5C and D**). With this approach, we found that the N-terminus of Aβ1-42 peptide had a higher density of half-tryptic sites, indicating increased structure-dependent digestion while being located within the Aβ fibril (**Fig. S5E**). Conversely, the mid-sequence and C-terminus were structurally less accessible for Proteinase K and more hidden within the Aβ fibril. Due to the different accessibility of Aβ1-42 peptide regions within the fibril, we included peptide pools deriving from the linear Aβ1-42 divided into N-terminal, mid-sequence, and C-terminal pools (**Table S4**). In addition, we included a peptide pool deriving from full-length 2N4R tau and axonal 0N4R tau variant representing both AD- and neurodegeneration-related antigens ^54^. For the tau pool, we selected linear peptides showing higher predicted affinity towards HLA molecules using the NetMHCIIpan database ^49,50^ (**Fig. S5F**). Moreover, tau-derived autoantigens have already been described in the general population, therefore we also tested tau peptides, which have previously been shown to have the highest percentage of total T cell responses ^55^ (**Table S5**). All selected peptides were 15-mers, which are preferentially presented by HLA class II to CD4^+^ T cells.

**Fig. 4:**
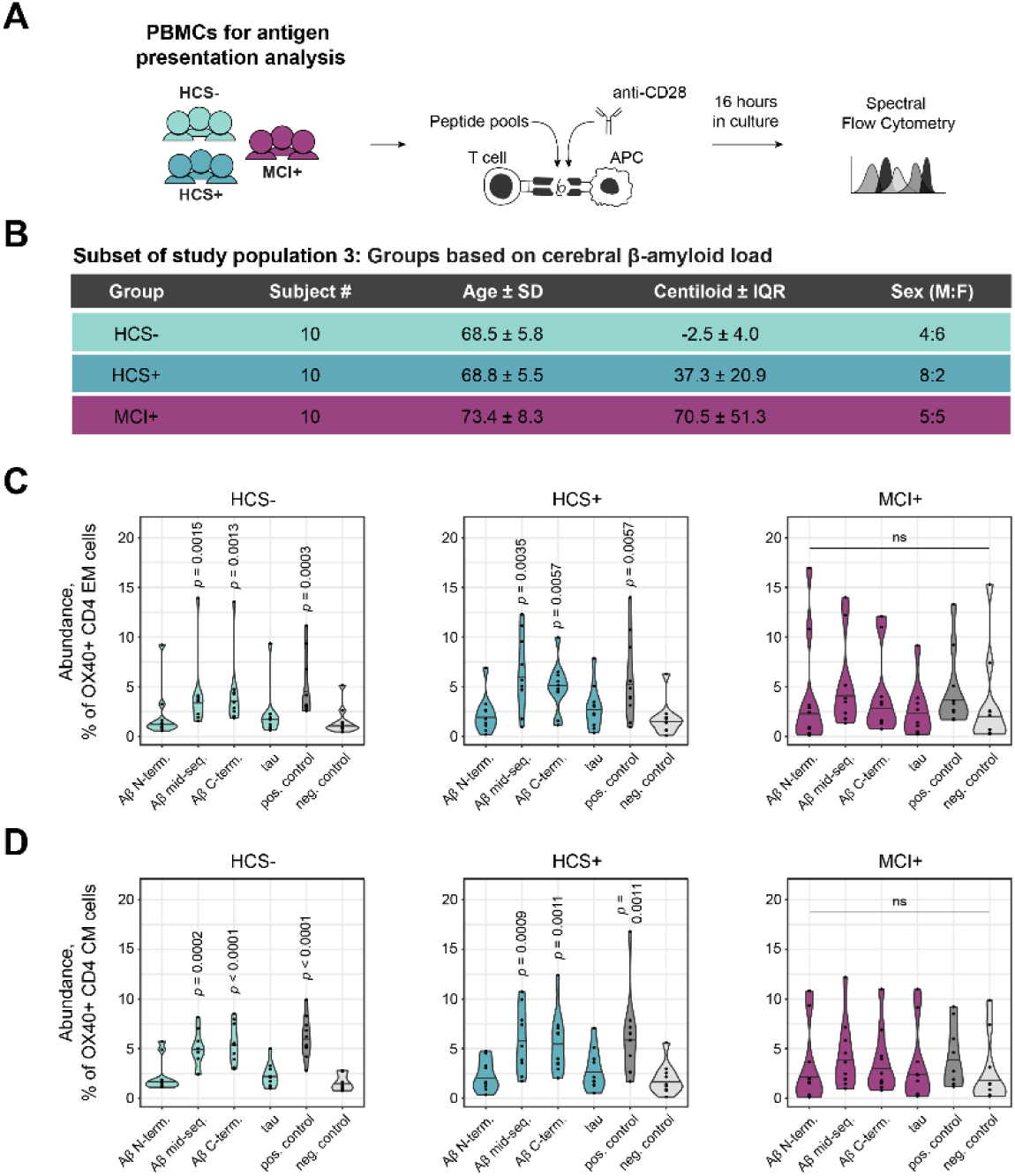
Altered T cell reactivity towards AD-related epitopes in subjects with early AD. (**A**) PBMCs from a subset of 30 subjects from study population 3 were pulsed for 16 hours with anti-CD28 and AD-related peptide pools. Expression of antigen-specific T cell markers were measured via spectral flow cytometry. (**B**) Demographic and clinical characteristics of study subset grouped according to cerebral Aβ load into Aβ-negative and -positive cognitively healthy subjects (HCS) and MCI subjects. (**C and D**) Violin plots with pooled data from the respective groups depicting the relative abundance of OX40^+^ cells among total CD4^+^ EM (**C**), and total CD4^+^ CM (**D**) T cells, upon pulsing with AD-related peptide pools. The *p*-values indicate statistical significance by comparison with unpulsed negative control. Statistical significance was calculated using non-parametric Kruskal-Wallis test with false-discovery-rate (FDR) method of Benjamini and Hochberg. Selected *p* values ≤ 0.1 (FDR 10%) are displayed.

We tested our antigen candidates with a subset of study population 3 that contained cerebral Aβ-negative and -positive HCS and MCI subjects (**Fig. 4B**). First, we compared positive control antigens with unpulsed conditions and selected optimal readout parameters for CD4^+^ EM (OX40) and CM (41BB, CD25, and OX40) T cells (**Fig. S6A-E**). Compared to the unpulsed condition, we identified autoreactive CD4^+^ EM and CM T cells responsive to Aβ mid-sequence and C-terminus, but not to N-terminal pools in HCS- and HCS+ subjects (**Fig. 4C and D**, **Fig. S6F and G**). These results stood in contrast to the LiP findings, which indicated that the Aβ N-terminus is most accessible for antigen processing and presentation, suggesting the involvement of immune tolerance towards the Aβ N-terminus. In addition, CD4^+^ EM and CM T cells of MCI+ subjects presented with a generally decreased reactivity to every peptide pool tested, including positive control antigens (**Fig. 4C and D**, **Fig. S6F and G**).

In summary, we report that compared to MCI, the preclinical AD stage (HCS+) is characterized by T cells with a less pro-inflammatory but more T_H_2 response-related immune profile. Moreover, CD4^+^ memory T cells showed an intact antigen-specific response to Aβ- derived linear peptide epitopes only in cognitively healthy subjects, but less so in subjects with MCI due to AD. Specifically, this immune response is directed against the mid-sequence and C-terminal fragments of Aβ peptide. This combined immunological framework could theoretically favour a protective adaptive immune response against pathological compounds in preclinical AD.

## Discussion

This study provides an in-depth analysis of the adaptive immune system in early AD by using state-of-the-art techniques, such as multidimensional mass cytometry combined with gene expression analysis via scRNA-seq, as well as single-cell secretome analysis and antigen presentation assays. Moreover, the study populations from the early AD spectrum used here are extensively characterized in terms of neuropsychological assessment, cerebral Aβ status, blood-based AD biomarkers, and anti-viral antibody titers. To our knowledge, this is therefore the first study characterizing immune cells of pre-symptomatic and early AD stages that incorporates modern analysis techniques with a comprehensive clinical assessment.

With this work, we demonstrate that peripheral and cerebral adaptive immune cell changes are functionally distinct in individuals in the early stages of AD. These early stages include preclinical AD (Aβ-positive and cognitively healthy) and prodromal AD, also referred to as MCI due to AD (Aβ-positive MCI). For the CD8^+^ cytotoxic subtype of T cells, we found substantially increased frequencies of highly antigen-specific TEMRA/effector cells in the blood of MCI cases and even in the CSF of preclinical AD cases. Moreover, these cells proved to have a less pro-inflammatory transcriptomic signature in preclinical AD, which was completely reversed in subjects in a more advanced disease stage, i.e. MCI due to AD. On a functional level, we conclude that CD8^+^ T cells in the blood of preclinical AD and MCI cases are expanded, but their target antigens still remain elusive. A different picture emerged for the CD4^+^ T helper and Treg subtype. Here, we observed a less immunosuppressive gene profile for Tregs in preclinical AD and MCI, leaving still the possibility of successfully mounting beneficial adaptive immune responses in preclinical AD. And indeed, functional analyses show that CD4^+^ memory T cells from preclinical AD subjects display a more T_H_2- like phenotype, secrete B cell-stimulating cytokines such as IL-4, and react against Aβ- derived antigens *in vitro*. In particular, our study demonstrates the presence of autoreactive CD4^+^ memory T cells preferentially targeting mid-sequence and C-terminus regions of the Aβ1-42 peptide. However, the more advanced MCI disease stage appeared to be characterized by less immunoreactivity, as any antigen-specific immune response previously seen in the cognitively healthy stage was no longer evident. Importantly, we show that our immunophenotyping results are independent of latent background infections with widespread *Herpesviridae*.

In line with our previous studies ^16^, we observed an increased abundance of CD8^+^ TEMRA/effector cells in peripheral blood and CSF of individuals in the early AD continuum. Similar results have previously been reported for CSF-derived cells from MCI and AD patients ^13^. As for other neurodegenerative diseases, increased numbers and clonal expansion of CD8^+^ TEMRA cells have also been observed in amyotrophic lateral sclerosis type 4 (ALS4) patients, suggesting their general involvement in neurodegeneration ^56^. In our study, the increased expression of pro-inflammatory gene modules in CD8^+^ TEMRA/effector cells of MCI subjects in combination with less immunosuppressive Treg function might indicate a contribution of CD8^+^ TEMRA/effector cells to the general hyper-inflammatory environment observed in MCI and AD. Indeed, a study using *in vitro* 3D modelling of the human brain showed increased infiltration of CD8^+^ T cells into AD brain cultures accompanied by exacerbated neuroinflammation and neurodegeneration ^57^. Compared to β- amyloid pathology, accumulation of neurofibrillary tangles made of hyperphosphorylated tau is more associated with neurodegeneration and cognitive decline ^58^. In line with this, in a previous study we also found increased numbers of CD8^+^ TEMRA/effector cells in individuals with high plasma levels of p-tau181 ^16^. This finding was replicated in the present study in MCI subjects with elevated plasma levels of p-tau217, an AD biomarker that correlates more strongly with brain Aβ and paired helical filament tau (PHFtau) tangles than p-tau181 ^59^. Moreover, recent studies suggest the involvement of CD8^+^ T cells in tau pathology, potentially driving neurodegeneration in response to tau-derived antigens ^60,61^. To better associate CD8^+^ TEMRA/effector immunophenotyping data with tau pathology, future studies and follow-up assessments will have to include more specific biomarkers tracking tau aggregation such as tau-PET or fluid biomarker candidates such as MTBR-tau243 ^62^. In future studies, we will design peptide pools deriving from the major components of AD pathology, Aβ and tau, and other sources that are preferentially presented to CD8^+^ T cells. So far, we have not found specific responses of CD8^+^ TEMRA/effector cells against AD- related (auto)antigens, but antigen presentation setups optimized for CD8^+^ T cells will address this matter. Importantly, one might have to consider also other CD8^+^ T cell subtypes that might be involved in AD pathology, as for example potentially beneficial, suppressive CD8^+^ PD-1^+^ T cells that might help shut down hyper-inflammatory microglia ^63^. In most cases, cytotoxic CD8^+^ T cells are known to react against cytoplasm-derived antigens such as viral or tumor antigens. CD8^+^ T cells reacting against viral epitopes have been detected in ALS, Parkinson’s Disease (PD), and AD patients ^13,56,64^. Virus infections could trigger the activation of cross-reactive T cells via molecular mimicry and antigen-independent bystander activation ^65^, and indeed the existence of cross-reactive T cells has been suggested in multiple sclerosis and narcolepsy ^66-68^.

For CD4^+^ T cells, we observed an intact reactivity towards Aβ-derived epitopes located within the mid-sequence and C-terminus of Aβ1-42 peptide in cognitively healthy individuals with and without cerebral Aβ burden. Conversely, we observed no reactivity towards Aβ N-terminus peptide pool. Previous studies reported increased T cell reactivity against Aβ- derived antigens in AD patients and age-matched cognitively healthy subjects ^18,19^. However, these studies did not perform a detailed AD biomarker analysis, namely Aβ PET scan and AD blood-based biomarkers. The lack of reactivity towards N-terminal fragments of Aβ suggests that autoreactive T cells against the Aβ N-terminus might be subjected to peripheral tolerance. Originally, upon cleavage of transmembrane self-protein APP by α- and γ-secretase in the non-amyloidogenic pathway, Aβ N-terminus is released within the sAPPα fragment and Aβ mid-sequence/C-terminus is part of the p3 peptide, making both available for processing and antigen-presentation by APCs, and ultimately for induction of peripheral immune tolerance ^69^. In contrast, Aβ1-42 peptide is a product of the amyloidogenic pathway, and during oligomerization and fibril formation, the C-terminus is located within the internal and hidden part of the Aβ fibril, and thus less accessible for structure-dependent digestion by APCs ^70-72^. This could result in autoreactive T cells specific for the C-terminus of Aβ1-42 that escape tolerance during AD pathology. As the N-terminal segment is located on the external and more accessible part of the Aβ fibril, it is prone to structure-based digestion ^70^, as shown by the LiP results. Hence, epitopes residing in the N-terminal segment might be preferentially presented by APCs and continue to induce strong peripheral tolerance, which might prevent a successful T cell response against N-terminal Aβ1-42.

The topic of immune tolerance in AD has been the subject of controversial debates. On the one hand, there are voices calling for therapeutic approaches that can limit immunosuppression and thus enable the adaptive immune system to act more effectively against pathological features of AD. Breaking immune tolerance using approaches such as immune checkpoint blockade of PD-1 or PD-1 ligand (PD-L1) in β-amyloidosis and tauopathy mouse models was shown to cause a systemic immune response, recruiting monocyte-derived macrophages to the brain, and resulting in reduced AD-associated pathology ^73,74^. Interestingly, blockade of the immune checkpoint in these experiments resulted not only in mobilization of monocytes to the brain but also in additional recruitment of CD4^+^ Tregs ^60,75^, suggesting that targeting PD-1 also alleviates the suppression of Tregs. On the other hand, it has been postulated that in common autoimmune diseases, but also in AD, it may be necessary to enhance Treg expansion and immunomodulatory function, e.g. by *ex vivo* expansion or *in vivo* treatment with IL-2 ^76-79^. Ultimately, both approaches appear to result in increased immunomodulatory potential of Tregs, which would be optimal for containing hyperinflammation and potentially cytotoxic CD8^+^ T cell responses in later AD stages, but they do not solve the problem of apparently suppressed beneficial T cell responses in preclinical AD.

CD4^+^ memory T cells displayed increased reactivity to AD-related antigens only in cognitively healthy subjects, while MCI subjects showed a general decrease in T cell immunoreactivity. In line with these results, studies using mouse models of β-amyloidosis showed that infiltrating T cells in brains with advanced β-amyloid pathology produced less pro-inflammatory cytokines ^80,81^. Indeed, we observed lower frequencies of T helper cell subsets producing IL-4 and IL-13 in MCI due to AD, which could result in less stimulation of antibody-producing B cells ^82-84^ and therefore less immunoglobulins potentially opsonizing Aβ plaques for clearance. Considering that Aβ accumulation in the brain begins decades before the onset of AD-related symptoms ^2,85^, T cell hypo-responsiveness in MCI due to AD might also be caused by exhaustion due to chronic antigen exposure. We reported previously that antigen presentation by myeloid-derived phagocytes is impaired under β-amyloid stress *in vitro* and *ex vivo* in brain-derived APCs from β-amyloidosis mouse models^81^. Impaired myeloid-derived phagocyte function and Aβ clearance might be among the causes for dysfunctional antigen presentation and subsequent reduced T cell activation observed in AD patients and β-amyloidosis mouse models ^80,81,86^.

In light of generally increased numbers and the more pro-inflammatory gene expression profile of CD8^+^ TEMRA/effector cells in MCI due to AD, this may be further evidence that CD8^+^ T cells are involved in AD pathology. Without knowledge of the target antigen, it remains speculative whether their role is beneficial or harmful. However, their pro-inflammatory, cytotoxic nature and activity in later disease stages suggest that CD8^+^ T cells, particularly TEMRA/effector cells, may further drive and exacerbate neuronal loss if responsive to antigens associated with AD pathology and if they reach the CNS in their activated state ^57,60^. On the other hand, the immune setup in preclinical AD still appears to be favourable for potentially beneficial CD4^+^ T cells targeting Aβ, because peripheral immunosuppression is downregulated in general, and CD4^+^ memory T cells show antigen-specific responses. This could, in theory, favor clearance of pathological Aβ aggregates via B cell stimulation. However, since CD4^+^ T cell specificity is aimed at parts of Aβ that are more hidden within the aggregated fibril structure, this response might miss its target. In fact, most β-amyloid-targeting antibodies, which have been tested to successfully elicit plaque clearance, bind epitopes within or near the N-terminus of the Aβ peptide ^87-89^. In contrast, T cell responses against the N-terminus of Aβ appear to be suppressed. Thus, there could be an epitope mismatch that complicates proper CD4^+^ T cell stimulation in preclinical AD. Future studies focusing on Tregs and peripheral tolerance in preclinical AD might provide further insight into this matter.

It is important to mention that our antigen candidate pools were not exhaustive, and in addition, the selection of peptide epitope candidates was quite stringent for long proteins such as tau, selecting only peptides with high affinity for HLA molecules. Potential autoantigens, however, might have lower HLA affinity to escape immune tolerance mechanisms. Moreover, so far we have mainly considered linear peptide antigens without modifications. But especially tau, with its multitude of post-translational modifications, such as phosphorylations at multiple sites, might harbour more potential T cell epitopes.

This research sheds light on the complex changes of the immune system in early AD stages and implies a more differentiated view on potential immunomodulatory therapies in AD. While it might be important in preclinical AD to support beneficial CD4^+^ T cell immune responses, it might be necessary in later stages to block harmful T cell responses that otherwise aggravate neurodegeneration. To assess potentially protective immune responses over time, it will be crucial to conduct long-term longitudinal studies.

## Supplementary Figures

**Fig. S1:**
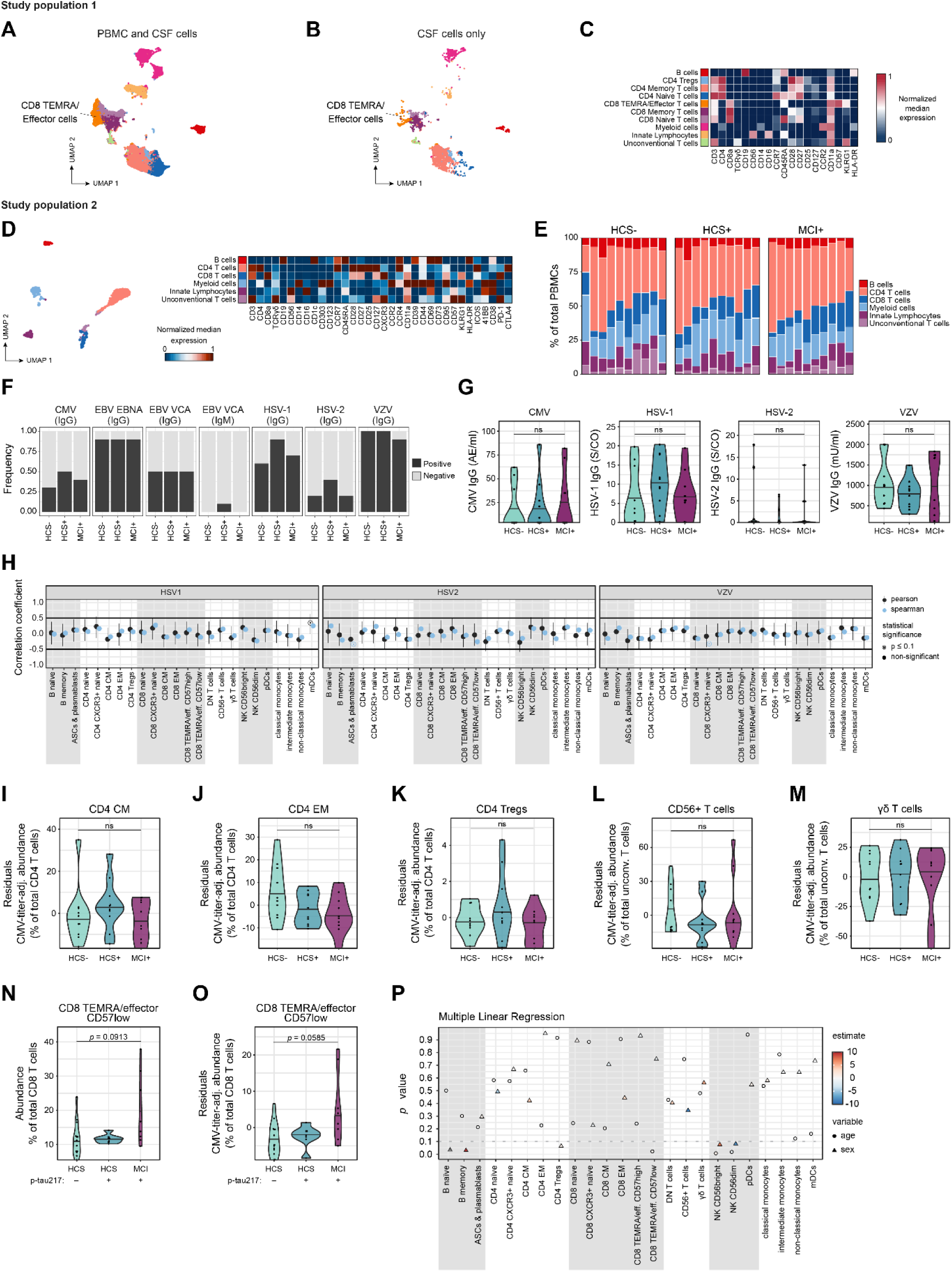
Immune composition and confounding factors in study populations 1 and 2. (**A and B**) **Study population 1:** UMAP plots depicting immune subsets based on paired FlowSOM clustering of PBMCs and CSF cells from study population 1. Depicted are immune subsets of PBMCs and CSF cells combined (**A**) and for CSF cells only (**B**). (**C**) Heatmap depicting median scaled marker expression for combined PBMC and CSF cell subsets. (**D**) **Study population 2:** UMAP coloured by PBMC subsets obtained with FlowSOM clustering. Corresponding heatmap depicts median scaled marker expression within PBMC subsets. (**E**) Proportions of PBMC subsets by diagnostic groups and individual subjects. (**F**) Proportions of study subjects positive for antiviral IgG or IgM titers against Cytomegalovirus (CMV), Epstein-Barr nuclear antigen (EBNA) of Epstein-Barr virus (EBV), viral capsid antigen (VCA) of EBV, Herpes simplex virus (HSV) 1 and 2, and Varicella-Zoster-Virus (VZV). (**G**) Violin plots showing quantitative *Herpesviridae* IgG titers across the 3 diagnostic groups. CMV in AE/ml (threshold ≥6), HSV-1 in S/CO (threshold >1), HSV-2 in S/CO (threshold >1), and VZV in mIU/ml (threshold >100). (**H**) Correlations of anti-*Herpesviridae* IgG titers with relative landmark node abundance. The estimated Pearson’s r correlation coefficients are shown in black, lines correspond to 95% confidence intervals; the estimated Spearman’s rho correlation coefficients are shown in blue. Empty circles indicate *p* values ≤ 0.1. (**I to M**) Violin plots showing relative abundance of landmark nodes CD4^+^ CM (**I**), CD4^+^ EM (**J**), and CD4^+^ Tregs (**K**) among total CD4^+^ T cells; as well as CD56^+^ T cells (**L**) and γδ T cells (**M**) among total unconventional T cells. (**N and O**) Alternative grouping of study population 2 according to the median of p-tau217 plasma levels (= 2.46 pg/ml). Violin plots showing relative abundance of CD8^+^ TEMRA/effector cells in p-tau217+ and p-tau217-subjects without (**N**) and with (**O**) adjustment for anti-CMV IgG titers. (**P**) Multiple linear regression for influence of age and sex on landmark node abundance. Statistical significance was calculated using non-parametric Kruskal-Wallis tests with false-discovery-rate (FDR) method of Benjamini and Hochberg.

**Fig. S2:**
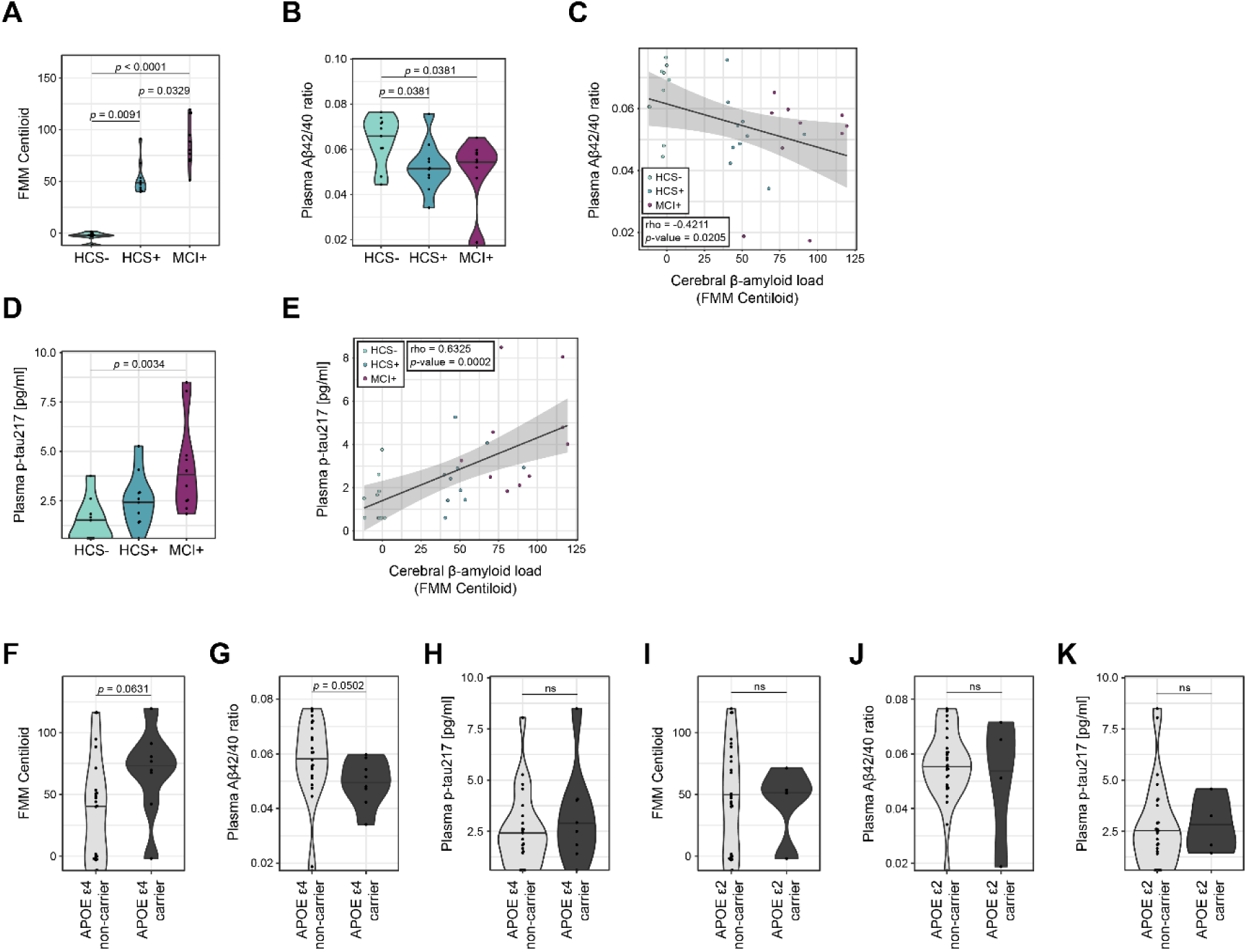
Demographic, clinical and genetic characteristics of study population 2. (**A and B**) Violin plots showing cerebral Aβ load (FMM Centiloid) (**A**) and plasma Aβ42/40 ratio (**B**) across the three diagnostic groups of study population 2. (**C**) Correlation of cerebral Aβ load (FMM Centiloid) with plasma Aβ42/40 ratio using linear regression. (**D**) Violin plots showing plasma p-tau217 levels across the three diagnostic groups of study population 2. (**E**) Correlation of cerebral Aβ load (FMM Centiloid) with plasma p-tau217 levels using linear regression. (**F to H**) Violin plots depicting cerebral Aβ load (FMM Centiloid) (**F**), plasma Aβ42/40 ratio (**G**), and plasma p-tau217 levels (**H**) in APOE-ε4 non-carriers and carriers. (**I to K**) Violin plots depicting cerebral Aβ load (FMM Centiloid) (**I**), plasma Aβ42/40 ratio (**J**), and plasma p-tau217 levels (**K**) in APOE-ε2 non-carriers and carriers. Statistical significance was calculated using non-parametric Kruskal-Wallis test with false-discovery-rate (FDR) method of Benjamini and Hochberg (**A, B and D**) or non-parametric unpaired Wilcoxon rank-sum test (**F to K**). Correlation coefficients (**C and E**) were calculated via Spearman approach (rho and *p* value reported). Grey shaded areas represent 95% confidence intervals of a linear regression. Selected *p* values ≤ 0.1 (FDR 10%) are displayed.

**Fig. S3:**
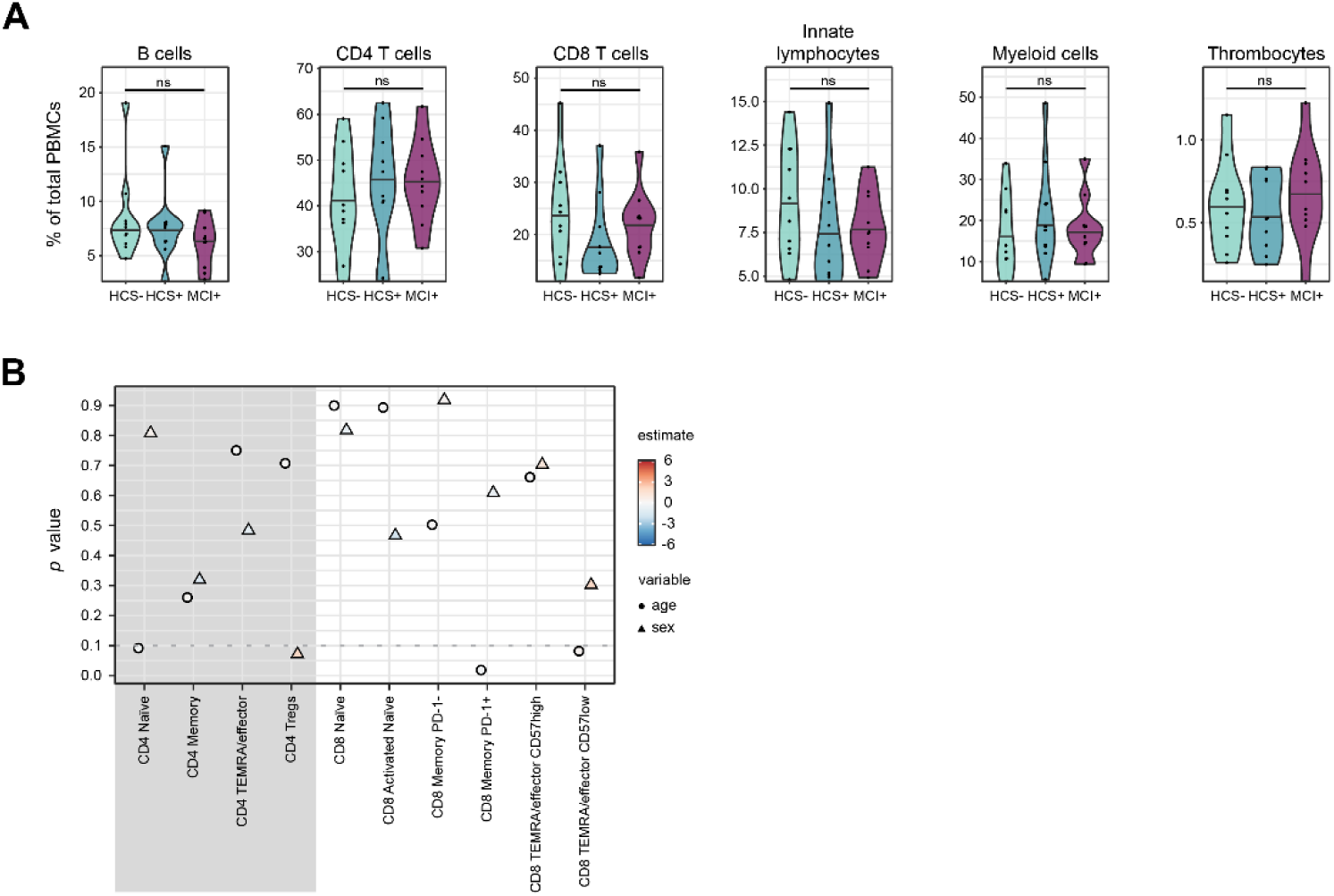
Single-cell transcriptomic analysis of peripheral immune cells of study population 2. (**A**) Violin plots showing relative abundance of main PBMCs subsets across the three diagnostic groups of study population 2. Statistical significance was calculated using non-parametric Kruskal-Wallis test with false-discovery-rate (FDR) method of Benjamini and Hochberg. (**B**) Multiple linear regression for influence of age and sex on CD4^+^ and CD8^+^ T cell cluster abundance.

**Fig. S4:**
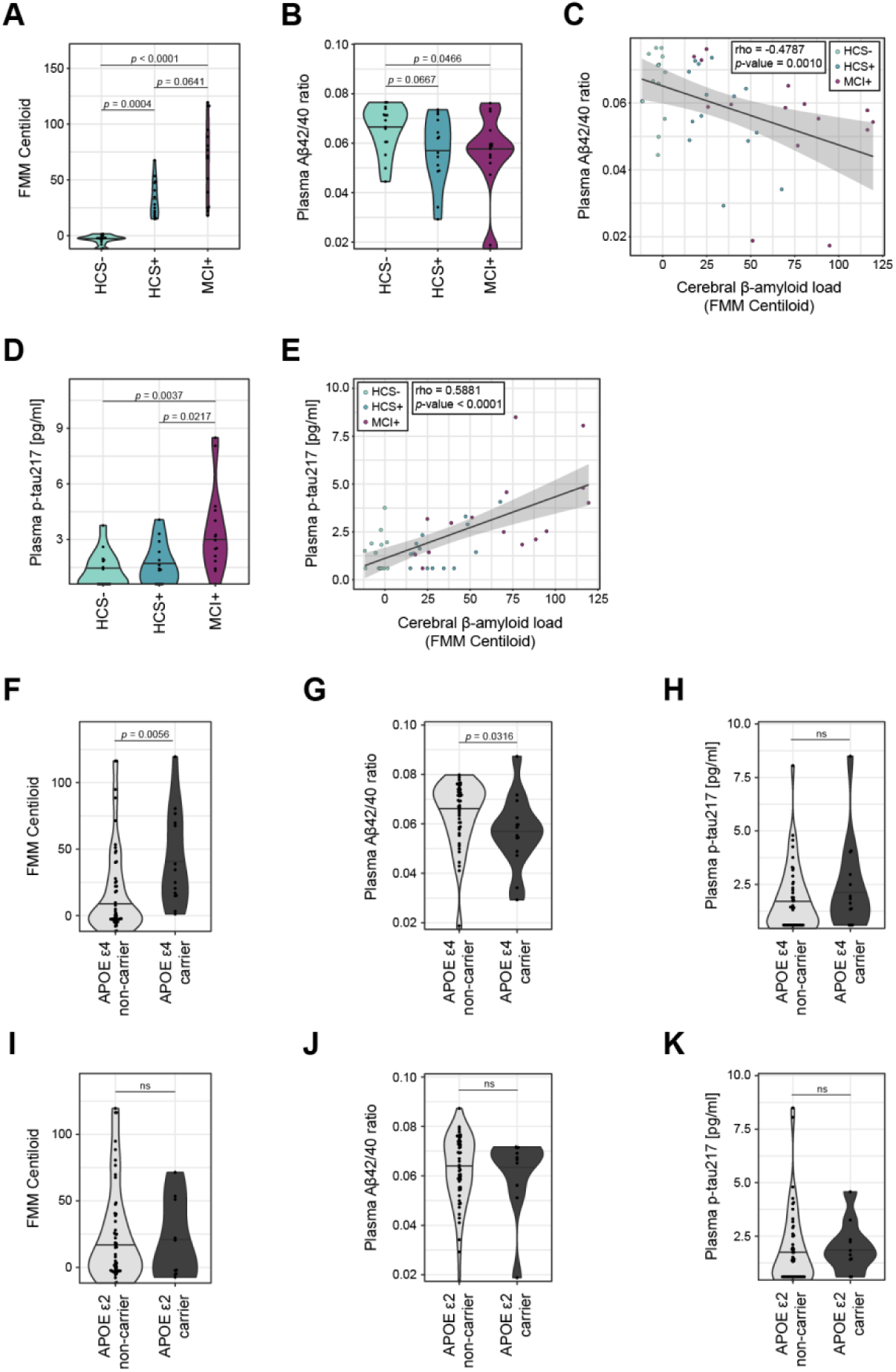
Characterization of study population 3. (**A and B**) Violin plots showing cerebral Aβ load (FMM Centiloid) (**A**) and plasma Aβ42/40 ratio (**B**) across the three diagnostic groups. (**C**) Correlation of cerebral Aβ load (FMM Centiloid) with plasma Aβ42/40 ratio using linear regression. (**D**) Violin plot showing plasma p-tau217 levels across the three diagnostic groups. (**E**) Correlation of cerebral Aβ load (FMM Centiloid) with plasma p-tau217 levels using linear regression. (**F to H**) Violin plots depicting cerebral Aβ load (FMM Centiloid) (**F**), plasma Aβ42/40 ratio (**G**), and plasma p-tau217 levels (**H**) in APOE-ε4 non-carriers and carriers. (**I to K**) Violin plots depicting cerebral Aβ load (FMM Centiloid) (**I**), plasma Aβ42/40 ratio (**J**), and plasma p-tau217 levels (**K**) in APOE-ε2 non-carriers and carriers. Statistical significance was calculated using non-parametric Kruskal-Wallis test with false-discovery-rate (FDR) method of Benjamini and Hochberg (**A, B and D**) or non-parametric unpaired Wilcoxon rank-sum test (**F to K**). Selected *p* values ≤ 0.1 (FDR 10%) are displayed. Correlation coefficients (**C and E**) were calculated via Spearman approach (rho and *p* value reported). Grey shaded areas represent 95% confidence intervals of a linear regression.

**Fig. S5:**
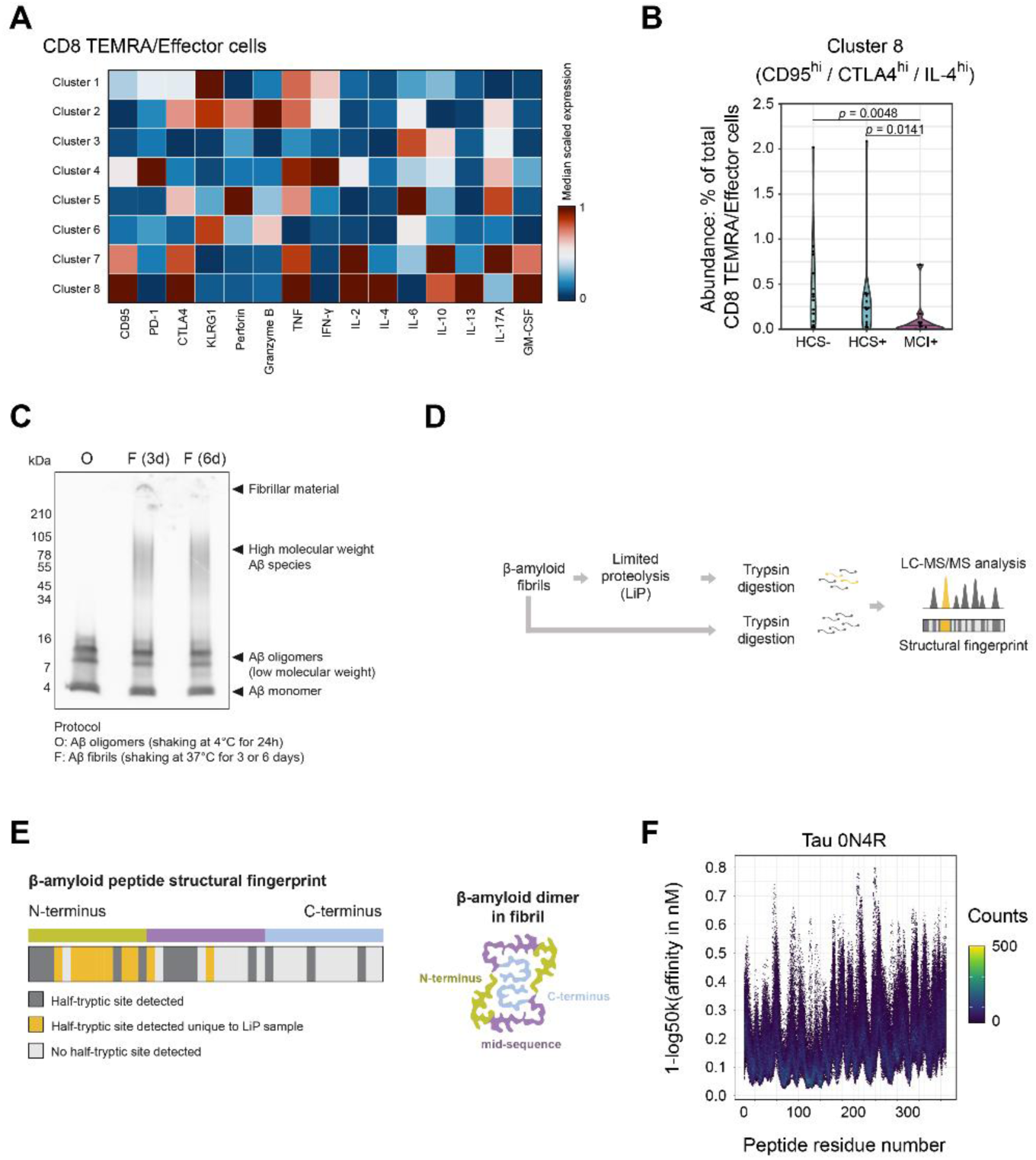
Secretome analysis for CD8^+^ TEMRA cells from study population 3 and sources of AD-related antigen candidates for antigen presentation analysis. (**A**) Heatmap depicting scaled median expression of eight functional clusters of CD8^+^ TEMRA/effector cells obtained via automated FlowSOM clustering using activation markers, cytokines, and cytotoxic granule content. (**B**) Violin plot showing relative abundance of Cluster 8 among total CD8^+^ TEMRA/effector cells. Statistical significance was calculated using non-parametric Kruskal-Wallis test with false-discovery-rate (FDR) method of Benjamini and Hochberg. Selected *p* values ≤ 0.1 (FDR 10%) are displayed. (**C**) Aβ peptide was aggregated *in vitro* for 24h at 4°C to obtain oligomers (= O) and 3 or 6 days at 37°C to produce high molecular weight oligomers, protofibrils and fibrils (= F). Resulting β-amyloid species were analyzed via SDS PAGE and Western Blot (protein load: 5μg/lane). Low molecular weight Aβ oligomers are found at approximately 9 kDa and 13.5 kDa, and high-molecular-weight Aβ aggregates are found above 30 kDa. (**D**) Workflow of limited proteolysis (LiP) for Aβ fibrils: Aβ fibrils are divided into a sample undergoing LiP and a control sample. Both samples are subject to tryptic digestion followed by LC-MS/MS and analysis of proteolytic fingerprints. (**E**) Left: barcode representing the structural fingerprint of Aβ1-42 peptide after LiP, from N- to C-terminus. Each residue from Aβ1-42 peptide is a vertical bar; residues with half-tryptic sites are shown in dark grey, residues with half-tryptic sites unique to the LiP sample are shown in yellow, and residues with no half-tryptic site are shown in light grey. Right: graphical representation of an Aβ dimer within fibril structure (top view). (**F**) Overlapping consecutive 15-mers derived from Tau 0N4R variant and their binding affinities (in nM) to HLA haplotypes included in the NetMHCIIpan database. Coloured by number of HLA haplotypes with similar binding affinity.

**Fig. S6:**
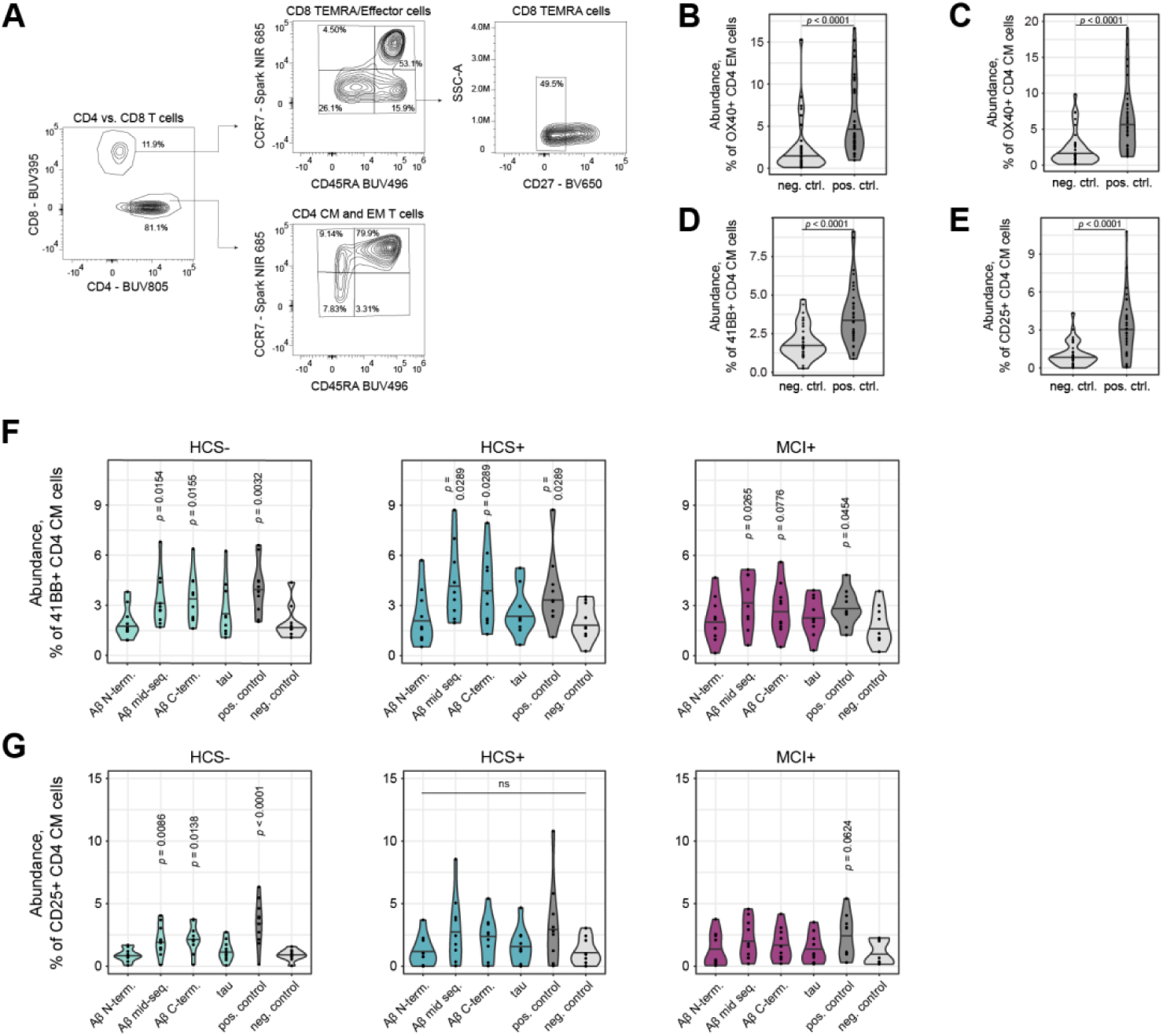
Antigen presentation analysis of subjects selected from study population 3. (**A**) Gating strategy to distinguish major CD4^+^ memory from CD8^+^ T cell subsets. (**B to E**) Violin plots with pooled data from all subjects depicting the relative abundance of OX40^+^ cells among total CD4^+^ EM cells (**B**), and the relative abundance of OX40^+^ (**C**), 41BB^+^ (**D**), and CD25^+^ (**E**) cells among total CD4^+^ CM cells upon pulsing with positive control peptide pools compared to negative control. (**F and G**) Violin plots with pooled data from subjects grouped according to cerebral Aβ load depicting the relative abundance of 41BB^+^ (**F**) and CD25^+^ (**G**) cells among total CD4^+^ CM cells upon pulsing with AD-related peptide pools. The *p* values in plots (**F and G**) indicate statistical significance by comparison with unpulsed negative control. Statistical significance was calculated using non-parametric unpaired Wilcoxon rank-sum test (**B to E**) or non-parametric Kruskal-Wallis test with false-discovery-rate (FDR) method of Benjamini and Hochberg (**F and G**). Selected *p* values ≤ 0.1 (FDR 10%) are displayed.

## Supplementary Tables

**Table S1:** Demographics and clinical characteristics of study population 1.

|  | Total | HCS- | HCS+ |
| --- | --- | --- | --- |
| Subjects | 9 | 5 | 4 |
| Age (mean ± SD) | 66.9 ± 8.6 | 65.4 ± 10.7 | 65.8 ± 7.8 |
| Sex (male/female) | 7/2 | 4/1 | 3/1 |
| APOE ε4 carriers | 55.6% | 60.0% | 50.0% |
| CSF Aβ40 (median ± IQR)<br>[pg/ml] | 8178.2 ± 3423.3 | 7262.0 ± 3175.0 | 9427.0 ± 3590.0 |
| CSF Aβ42 (median ± IQR)<br>[pg/ml] | 636.3 ± 241.8 | 627.0 ± 222.0 | 601.0 ± 248.0 |
| CSF Aβ42/ Aβ40<br>(median ± IQR) | 0.076 ± 0.013 | 0.086 ± 0.007 | 0.065 ± 0.004 |

**Table S2:** Demographics and clinical characteristics of study population 2.

|  | Total | HCS- | HCS+ | MCI+ |
| --- | --- | --- | --- | --- |
| Subjects | 30 | 10 | 10 | 10 |
| Age (mean ± SD) | 72.1 ± 6.5 | 69.6 ± 4.3 | 69.5 ± 6.9 | 77.3 ± 4.9 |
| Sex (male/female) | 15/15 | 5/5 | 7/3 | 3/7 |
| Years of education<br>(mean ± SD) | 14.8 ± 3.1 | 16.0 ± 3.2 | 15.8 ± 2.9 | 12.7 ± 1.8 |
| MMSE (mean ± SD) | 27.0 ± 1.9 | 29.7 ± 0.5 | 28.8 ± 1.8 | 27.0 ± 1.9 |
| APOE ε4 carriers | 26.7% | 10.0% | 30.0% | 40.0% |
| APOE ε2 carriers | 13.3% | 10.0% | 10.0% | 20.0% |
| FMM Centiloid (median ± IQR) | 47.8 ± 75.9 | -2.2 ± 2.1 | 47.8 ± 10.0 | 84.6 ± 38.2 |
| Plasma Aβ40 (median ± IQR)<br>[pg/ml] | 110.5 ± 31.9 | 115.5 ± 34.7 | 107.0 ± 19.7 | 109.0 ± 43.7 |
| Plasma Aβ42 (median ± IQR)<br>[pg/ml] | 5.8 ± 2.7 | 7.6 ± 1.6 | 5.5 ± 1.2 | 5.3 ± 1.4 |
| Plasma Aβ42/ Aβ40<br>(median ± IQR) | 0.056 ± 0.016 | 0.068 ± 0.011 | 0.051 ± 0.008 | 0.055 ± 0.010 |
| Plasma NfL (median ± IQR)<br>[pg/ml] | 27.0 ± 14.2 | 22.2 ± 9.5 | 27.1 ± 14.8 | 31.2 ± 5.9 |
| Plasma GFAP (median ± IQR) | 144.0 ± 91.0 | 126.0 ± 45.5 | 122.5 ± 75.0 | 198.5 ± 102.5 |
| [pg/ml] |  |  |  |  |
| Plasma p-tau217<br>(median ± IQR)<br>[pg/ml] | 2.5 ± 2.2 | 1.1 ± 1.2 | 2.5 ± 1.4 | 3.6 ± 2.2 |

**Table S3:** Demographics and clinical characteristics of study population 3.

|  | Total | HCS- | HCS+ | MCI+ |
| --- | --- | --- | --- | --- |
| Subjects | 59 | 15 | 14 | 15 |
| Age (mean ± SD) | 70.1 ± 8.2 | 67.0 ± 6.8 | 69.3 ± 6.5 | 75.3 ± 7.5 |
| Sex (male/female) | 33/26 | 4/11 | 12/2 | 6/9 |
| Years of education<br>(mean ± SD) | 14.7 ± 2.9 | 14.8 ± 3.0 | 16.1 ± 2.2 | 14.1 ± 3.1 |
| MMSE (mean ± SD) | 28.4 ± 1.6 | 29.5 ± 0.8 | 27.9 ± 1.5 | 27.5 ± 1.8 |
| APOE ε4 carriers | 29.5% | 6.7% | 50.0% | 33.3% |
| APOE ε2 carriers | 18.2% | 20% | 21.4% | 13.3% |
| FMM Centiloid (median ± IQR) | 9.7 ± 42.4 | -2.4 ± 3.3 | 26.4 ± 26.7 | 71.4 ± 59.4 |
| Plasma Aβ40 (median ± IQR)<br>[pg/ml] | 111.0 ± 33.6 | 114.0 ± 23.5 | 114.5 ± 33.4 | 102.0 ± 36.3 |
| Plasma Aβ42 (median ± IQR)<br>[pg/ml] | 7.0 ± 2.6 | 7.5 ± 1.3 | 6.0 ± 2.7 | 5.7 ± 2.7 |
| Plasma Aβ42/ Aβ40<br>(median ± IQR) | 0.064 ± 0.017 | 0.069 ± 0.012 | 0.059 ± 0.019 | 0.059 ± 0.009 |
| Plasma NfL (median ± IQR)<br>[pg/ml] | 20.2 ± 13.4 | 19.5 ± 9.4 | 18.4 ± 6.7 | 30.1 ± 10.3 |
| Plasma GFAP (median ± IQR)<br>[pg/ml] | 114.0 ± 69.8 | 108.0 ± 66.0 | 102.2 ± 37.4 | 195.0 ± 109.0 |
| Plasma p-tau217<br>(median ± IQR)<br>[pg/ml] | 1.4 ± 1.9 | 0.6 ± 1.3 | 1.4 ± 1.6 | 3.0 ± 2.3 |

**Table S4:**
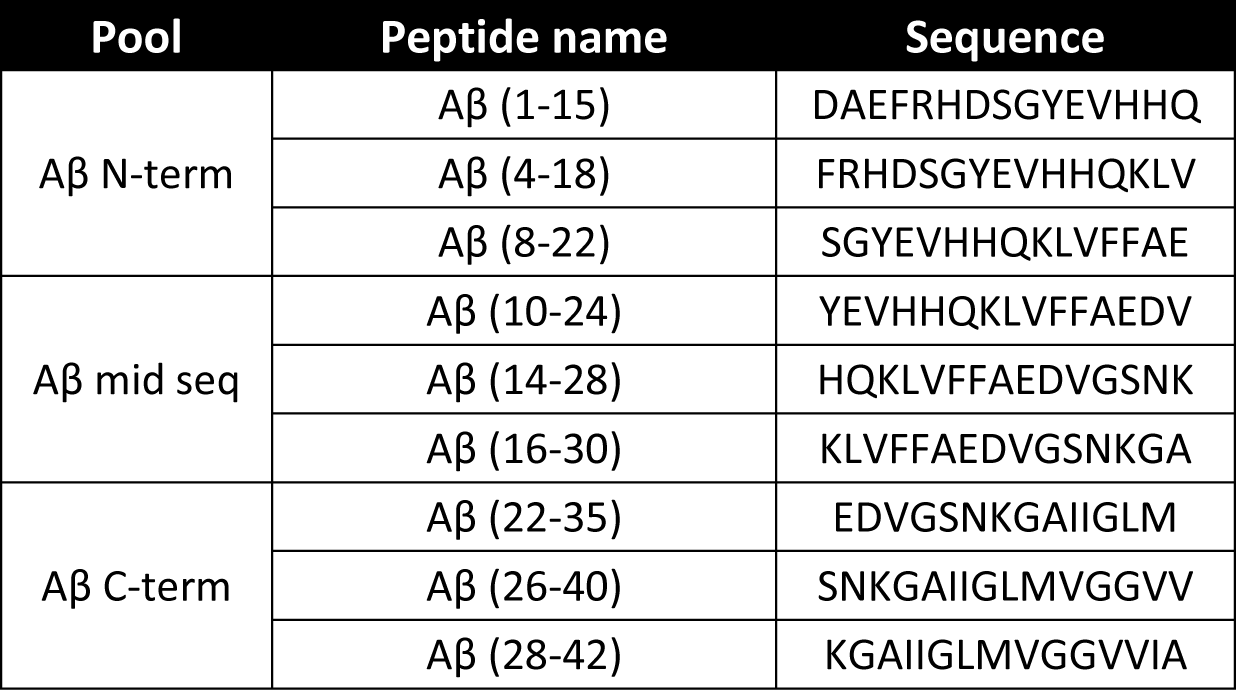
Linear Aβ1-42-derived epitopes used for T cell antigen presentation.

**Table S5:**
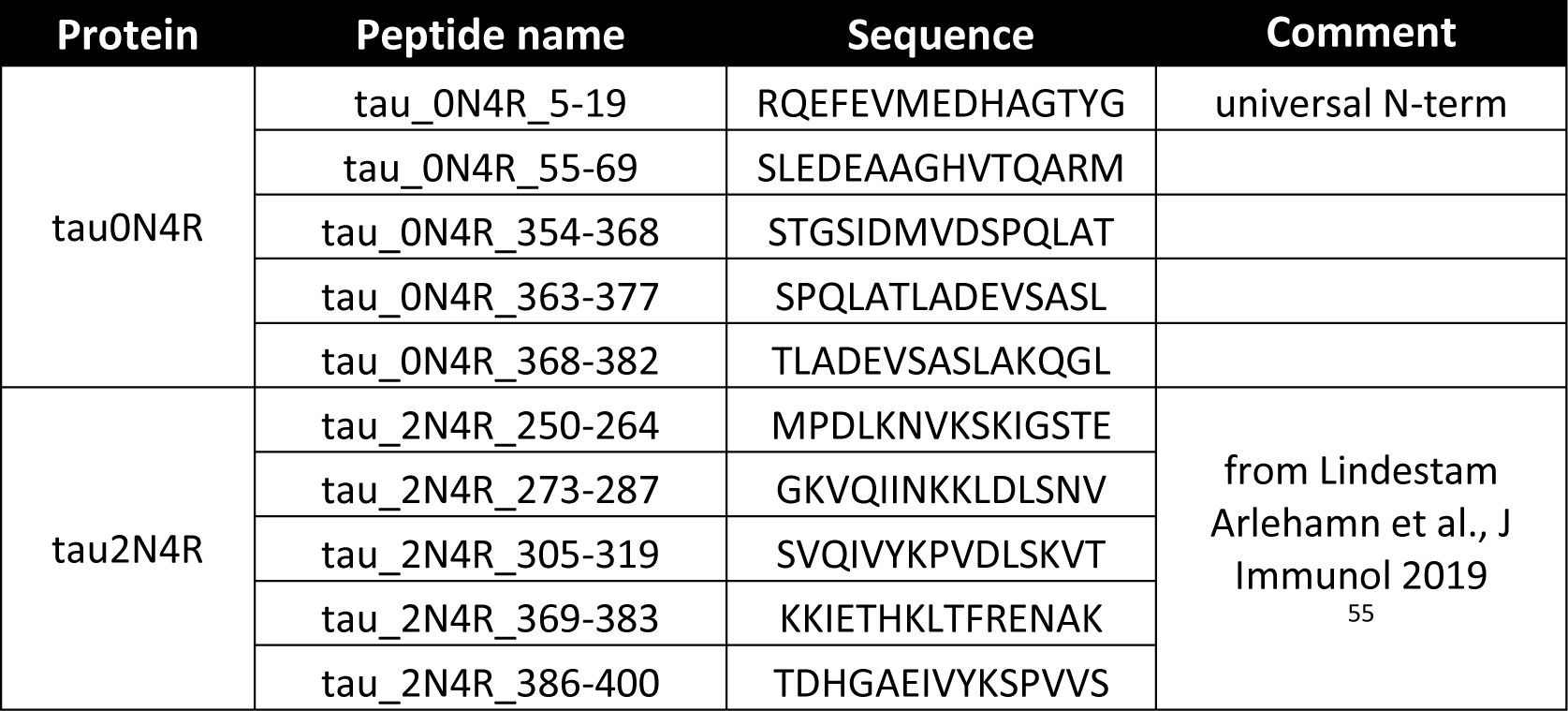
Linear tau-derived epitopes used for T cell antigen presentation.

**Table S6:** Heavy metal-labeled antibodies used for mass cytometry.

| Isotope | Metal | Antigen | Clone | Supplier | Category | Antibody class |
| --- | --- | --- | --- | --- | --- | --- |
| 89 | Y | CD45 | HI30 | Fluidigm | barcoding | na |
| 106 | Cd | CD45 | HI30 | Biolegend | barcoding | na |
| 110 | Cd | CD45 | HI30 | Biolegend | barcoding | na |
| 111 | Cd | CD45 | HI30 | Biolegend | barcoding | na |
| 112 | Cd | CD45 | HI30 | Biolegend | barcoding | na |
| 113 | In | CD45 | HI30 | Biolegend | barcoding | na |
| 114 | Cd | CD45 | HI30 | Biolegend | barcoding | na |
| 116 | Cd | CD8a | IA6-2 | Fluidigm | surface | type |
| 141 | Pr | CD45RA | HI100 | Biolegend | surface | type |
| 142 | Nd | CD19 | HIB19 | Fluidigm | surface | type |
| 143 | Nd | CD123 | 6H6 | Fluidigm | surface | type |
| 144 | Nd | CD69 | FN50 | Fluidigm | surface | state |
| 145 | Nd | CD4 | RPA-T4 | Fluidigm | surface | type |
| 146 | Nd | CD11a | HI111 | Biolegend | surface | state |
| 147 | Sm | CD303 | 201A | Fluidigm | surface | none |
| 148 | Nd | CD137 | 4B4-1 | Biolegend | surface | state |
| 149 | Sm | CD25 | 2A3 | Fluidigm | surface | state |
| 150 | Nd | CD127 | A019D5 | Biolegend | surface | state |
| 151 | Eu | CD14 | M5E2 | Fluidigm | surface | type |
| 152 | Sm | KLRG1 | 13F12F2 | ThermoFisher | surface | state |
| 153 | Eu | CD192 | K036C2 | Fluidigm | surface | state |
| 154 | Sm | CD1c | AD5-8E7 | Miltenyi Biotec | surface | type |
| 155 | Gd | CD279 | EH12.2H7 | Fluidigm | surface | state |
| 156 | Gd | CD183 | G025H7 | Fluidigm | surface | state |
| 158 | Gd | CD194 | L291H4 | Fluidigm | surface | state |
| 159 | Tb | CD197 | G043H7 | Fluidigm | surface | type |
| 160 | Gd | CD28 | CD28.2 | Fluidigm | surface | type |
| 161 | Dy | CD152 | 14D3 | Fluidigm | surface | state |
| 162 | Dy | TCRgd | B1 | Biolegend | surface | type |
| 163 | Dy | CD57 | HCD57 | Fluidigm | surface | state |
| 164 | Dy | CD95 | DX2 | Fluidigm | surface | state |
| 165 | Ho | CD39 | A1 | Biolegend | surface | state |
| 166 | Er | CD44 | BJ18 | Fluidigm | surface | state |
| 167 | Er | CD27 | L128 | Fluidigm | surface | type |
| 168 | Er | CD73 | AD2 | Fluidigm | surface | state |
| 169 | Tm | CD45 | HI30 | Fluidigm | barcoding | na |
| 170 | Er | CD3 | UCHT1 | Fluidigm | surface | type |
| 171 | Yb | CD45 | HI30 | Fluidigm | barcoding | na |
| 172 | Yb | CD38 | HIT2 | Fluidigm | surface | state |
| 173 | Yb | CD45 | HI30 | Fluidigm | barcoding | none |
| 174 | Yb | HLA-DR | L243 | Fluidigm | surface | state |
| 175 | Lu | ICOS | C398.4A | Fluidigm | surface | state |
| 176 | Yb | CD56 | NCAM16.2 | Fluidigm | surface | type |
| 198 | Pt | Live/dead | - | Fluidigm | live cell<br>discrimination | none |
| 209 | Bi | CD16 | 3G8 | Fluidigm | surface | type |

**Table S7:** Antibody panel used for T cell secretome and antigen-presentation experiments.

| Fluorochrome | Antigen | Clone | Supplier | Panel |
| --- | --- | --- | --- | --- |
| - | L/D blue | - | Invitrogen, L34961 | secretome + antigen-presentation |
| Pacific Blue | CD3 | UCHT1 | Biolegend | secretome + antigen-presentation |
| BUV395 | CD8 | RPA-T8 | BD, 563796 | secretome + antigen-presentation |
| BUV805 | CD4 | RPA-T4 | BD, 742000 | secretome + antigen-presentation |
| BV605 | CD56 | 5.1H11 | Biolegend, 362537 | secretome + antigen-presentation |
| KIRAVIA Blue | FoxP3 | 206D | Biolegend, 320131 | secretome + antigen-presentation |
| PE/Fire 700 | TCRgd | B1 | Biolegend, 331237 | secretome + antigen-presentation |
| BUV 496 | CD45RA | 206D | BD, 741182 | secretome + antigen-presentation |
| Spark NIR 685 | CCR7 | G043H7 | Biolegend, 353257 | secretome + antigen-presentation |
| BV650 | CD27 | O323 | Biolegend, 302827 | secretome + antigen-presentation |
| BV510 | CD57 | QA17A04 | Biolegend, 393313 | secretome + antigen-presentation |
| APC/Fire 810 | CXCR5 | J252D4 | Biolegend, 356955 | secretome + antigen-presentation |
| PE/Fire 810 | KLRG1 |  | Biolegend, 367733 | secretome + antigen-presentation |
| PE/Fire 640 | CD95 | DX2 | Biolegend, 305657 | secretome |
| PE/Cy5 | PD1 | EH12.2H7 | Biolegend, 329971 | secretome |
| BB515 | CD25 | BC96 | BD, 564468 | secretome + antigen-presentation |
| BB700 | CTLA4 | BNI3 | BD, 566902 | secretome |
| BV 711 | TNF | MAb11 | Biolegend, 502939 | secretome + antigen-presentation |
| AF 700 | GZMB | QA16A02 | Biolegend, 372221 | secretome + antigen-presentation |
| APC/Cy7 | Perforin | dG9 | Biolegend, 308127 | secretome + antigen-presentation |
| BV 750 | IFN-γ | 4S.B3 | Biolegend, 502549 | secretome + antigen-presentation |
| BUV 737 | IL-2 | MQ1-17H12 | BD, 612836 | secretome + antigen-presentation |
| PE-Cy7 | IL-4 | MP4-25D2 | Biolegend, 500823 | secretome + antigen-presentation |
| PE/Dazzle 594 | IL-6 | MQ2-13A5 | Biolegend, 501121 | secretome |
| AF 647 | IL-10 | JES3-9D7 | Biolegend, 501414 | secretome |
| BV421 | IL-13 | JES10-5A2 | Biolegend, 501915 | secretome + antigen-presentation |
| BV785 | IL-17A | BL168 | Biolegend, 512337 | secretome + antigen-presentation |
| PE | GM-CSF | BVD2-21C11 | Biolegend, 502305 | secretome + antigen-presentation |
| PE-Dazzle 594 | 41BB | 4B4-1 | Biolegend, 309825 | antigen-presentation |
| AF 647 | OX40 | Ber-ACT35 | Biolegend, 350017 | antigen-presentation |
| PE/Cy5 | CD154 | 24-31 | Biolegend, 310808 | antigen-presentation |

## Acknowledgements

We would like to thank all the volunteers who kindly participated in the studies. We thank all study physicians and neuropsychologists for their contribution to the assessments and conduct of the study. Samples from participants were collected at the Center for Prevention and Dementia Therapy (Institute for Regenerative Medicine, University of Zurich) by study nurses led by Esmeralda Gruber. We would like to thank the Cytometry Facility (University of Zurich) for technical assistance, and the Institute of Medical Virology (University of Zurich) for virus serology analysis. Special thanks go to Prof. Paola Picotti from the Institute of Molecular Systems Biology (ETH Zurich) for providing expertise on limited proteolysis and mass spectrometry. Finally, we would like to thank Hayder Shweliyya and his team at the Sahlgrenska University Hospital for fluid biomarker analyses.

## Author contributions

Conceptualization, C.R., V.T., A.G., and C.G.; Methodology, C.R., A.M., L.H., L.F., S.S., C.S., T.K., L.K., K.B., H.Z., and C.G.; Software, C.R. and A.M.; Formal Analysis, C.R. and C.G.; Investigation, C.R. and C.G.; Resources, C.H., V.T., and A.G.; Data Curation, C.R. and A.M.; Writing - Original Draft, C.R. and C.G.; Writing - Review & Editing, A.M., C.S., K.B., H.Z., M.T.F., L.K., V.T., A.G., and C.G.; Visualization, C.R.; Supervision and Project Administration, V.T., A.G., and C.G.; Funding Acquisition, C.R., K.B., H.Z., C.H., R.M.N., A.G., and C.G.

## Consent statement

All study participants gave written informed consent. All studies and further use for the current analyses were approved by the local ethics committee (Kantonale Ethikkommission Zürich) and conducted in accordance with their guidelines and the Declaration of Helsinki.

